# Reciprocal Fronto-Parietal Interactions Support Distinct Motor Strategies during Sequential Reaching

**DOI:** 10.64898/2026.03.03.709252

**Authors:** Georgios Bardanikas, Shrabasti Jana, Nicolas Meirhaeghe, Frederic Barthelemy, Alexa Riehle, Sonja Grün, Andrea Brovelli, Thomas Brochier

## Abstract

Motor planning is a highly flexible process that adapts to changing behavioral demands. Frontal and parietal circuits are considered as critical nodes that support planning processes. However, it remains unclear whether the coordination of motor and parietal areas is preserved or adjusted in response to varying behavioral contexts. To address this open question, we trained two rhesus macaques to perform a visually-guided sequential reaching task, in which they could adapt their behavior to varying degrees of target predictability within the sequence. Analysis of eye and hand movements revealed distinct visuomotor strategies that were reflected in corresponding neural activity patterns. During motor anticipation, the direction of the upcoming reach toward predictable targets could be decoded from preparatory neural activity prior to target onset both in motor regions (PMd/M1) and parietal area 7A, whereas during reactive movements directional information emerged only after target onset. Using feature-specific information transfer analysis, we found that information about the upcoming movement direction was transmitted between 7A and PMd/M1 through bidirectional interactions. This pattern of interactions was preserved during motor anticipation and shifted earlier in time with respect to target onset. Our findings thus support a reciprocal fronto-parietal network that flexibly adjusts the timing of preparatory activity as different strategies are employed under varying behavioral constraints. Contrary to classical hierarchical models predicting serial activation across parietal and motor areas, parietal-to-motor interactions did not precede motor-to-parietal interactions, consistent with a heterarchical organization of cortical processing during flexible motor planning.

## Introduction

Motor planning is widely recognized as the process by which the motor commands required to achieve a selected goal is prepared before movement initiation (Wong et al., 2015). Far from being a mere reflexive response to visual target presentation, this process demonstrates remarkable adaptability to environmental contexts. Previous studies have shown that motor planning can optimize movements toward targets with varying spatial predictability (Bastian et al., 2003), intercept moving targets by anticipating their trajectory (Van Donkelaar et al., 1992; Merchant & Georgopoulos, 2006), or plan entire movement sequences in advance (Shima & Tanji, 2000; Averbeck et al., 2002, 2003; Lu & Ashe, 2005). However, natural situations often involve sequences of movements where information about upcoming actions is dynamic or incomplete, necessitating real-time adjustments in motor strategies. For example during a long tennis rally, a player must not only prepare a shot based on the incoming ball’s trajectory, but also anticipate the next moves by accounting for the opponent’s positioning and playing style. Here, we investigate how motor planning flexibly adapts across different motor strategies during sequential reaching.

At the neural level, motor planning relies on the coordinated activity of distributed cortical regions involved in visuomotor transformation. The posterior parietal cortex (PPC) plays a central role in integrating sensory and contextual information to estimate future sensory states, and represent movement intentions and spatial goals (Constantinidis & Steinmetz, 2001; Andersen & Buneo, 2002; Blakemore & Sirigu, 2003; Cui & Andersen, 2007; Fontana et al., 2012; Krause et al., 2014). Within PPC, area 7A has been closely linked to upcoming eye and arm movements (Hyvärinen & Poranen, 1974; MacKay, 1992; Heider et al., 2010). However, several neurophysiological studies have shown that 7A neurons do not only encode low-level visual features, but also higher-order cognitive processes, such as maze solution (Crowe et al., 2004a, 2005), and planned movement direction before interception of moving targets (Li et al., 2022). Recurrent population dynamics in 7A have been proposed to estimate sensory states through an internal model, providing a mechanism for encoding prior information and updating predictions (Lakshminarasimhan et al., 2023). Mounting evidence thus suggests that area 7A is involved in higher order cognitive and planning processes.

Visuomotor information represented in 7A must ultimately be transformed into motor commands. Premotor cortex contributes to this process by selecting the final action and encode movement parameters before movement onset, thereby reflecting motor preparation (Georgopoulos et al., 1982, 1986; Riehle & Requin, 1989; Crammond & Kalaska, 2000; Cisek & Kalaska, 2005; Thura & Cisek, 2014; Churchland & Shenoy, 2024). Anatomical studies have revealed reciprocal axonal projections between area 7A and premotor cortex (Markov et al., 2014; Chaudhuri et al., 2015), suggesting that sensory representations in 7A can influence motor planning. From a dynamical systems perspective, preparatory activity is thought to establish the initial neural state from which movement-related dynamics unfold (Shenoy et al., 2013; Gallego et al., 2017; Vyas et al., 2020). Parietal signals encoding predicted sensory states (Lakshminarasimhan et al., 2023) can then bias motor dynamics toward these initial conditions earlier, facilitating faster and more efficient motor planning and execution (Churchland et al., 2006, 2010, 2012). Together, converging evidence suggests that premotor regions and the posterior parietal area 7A represent critical hubs supporting flexible motor planning during goal-directed actions. However, it remains unclear whether the coordination of motor and parietal areas is preserved or adjusted as different motor strategies are employed to meet context-dependent demands.

Hierarchical models of cortical processing provide testable predictions about characteristic delays between the serial activations of cortical regions (Felleman & Van Essen, 1991; Shipp, 2005; Fries, 2005; Siegel et al., 2015; Vinck et al., 2023). Based on these models, parietal areas should activate prior to motor areas. Alternative frameworks suggest that during complex behavior, cortical processing is distributed across multiple interacting areas with reciprocal interactions between parietal and motor areas (Steinmetz et al., 2019; Findling et al., 2025; Wang et al., 2025; International Brain Laboratory et al., 2025; Rosen & Freedman, 2025).

To test these hypotheses, we simultaneously recorded neural activity from the dorsal premotor and primary motor cortex (PMd/M1) and the posterior parietal area 7A in macaque monkeys performing a sequential reaching task with increasing target predictability. Population-level dynamics and interareal interactions were studied using mesoscopic multi-unit activity (MUA). We observed that monkeys adopted two different behavioral strategies to adapt to behavioral context: one monkey mainly anticipated reaches to predictable targets, whereas the other reacted at similar latency irrespective of target predictability. These differences were reflected in distinct neural activity patterns, with neural representations of upcoming movement direction emerging before or after target onset depending on the employed behavioral strategy. We found that information about the upcoming movement direction was contained in bidirectional, rather than serial, interactions between PMd/M1 and 7A. Such bidirectional interactions were temporally advanced in the monkey showing motor anticipation. Together, these findings indicate that parieto-motor communication is dynamically organized according to behavioral context, rather than reflecting a fixed serial flow from parietal to motor regions. Within this framework, reciprocal interactions between area 7A and PMd/M1 may provide a flexible neural substrate for integrating predictive and reactive control during sequential motor behavior.

## Results

### Experimental task design

Two macaque monkeys (J & E) performed a visually-guided sequential reaching task in which target predictability was systematically manipulated to promote motor anticipation within the sequence. Monkeys were sitting in a Kinarm primate chair (Bkin technologies), and were trained to perform movements with their right arm attached to an articulated robotic exoskeleton controlling a cursor on a 2D horizontal screen in front of them (**Figure 1A**). Starting from a unique central location, monkeys were required to reach 3 visual targets presented sequentially at one of 6 possible peripheral locations (Jana et al., 2025). After each target presentation, with no instructed preparatory delay, monkeys controlled the exoskeleton to reach the Valid Target Area (defined as a circle with a radius of 1 cm around the target, see Methods) and to keep it within its limit for a brief 200 ms delay (**Figure 1B**). Monkeys were rewarded after having correctly reached three targets presented in sequence (three examples in **Figure 1C**). At each trial, one out of 12 predefined 3-target sequences was pseudo-randomly selected (**Figure S.1**). Importantly, the 12 sequences were designed in such a way that the location of the target becomes more predictable with increasing target rank in the sequence (examples in **Figure 1D**). Specifically, the first target was presented at 1 of the 6 possible positions (distributed over 360° around the central target) and required a center-out reaching movement. The second target was presented at 1 of 2 possible positions and the third target position was fully determined by the two previous ones. Both the 2^nd^ and 3^rd^ targets required hand movements starting and ending at a peripheral target. Thus, this task allowed the comparison of arm movements (and the underlying neural activity) across two different experimental conditions; when the predictability of the forthcoming target location and hence the possibility for movement anticipation was absent (1^st^ target) or gradually higher (2^nd^ and 3^rd^ targets).

**Figure 1.**
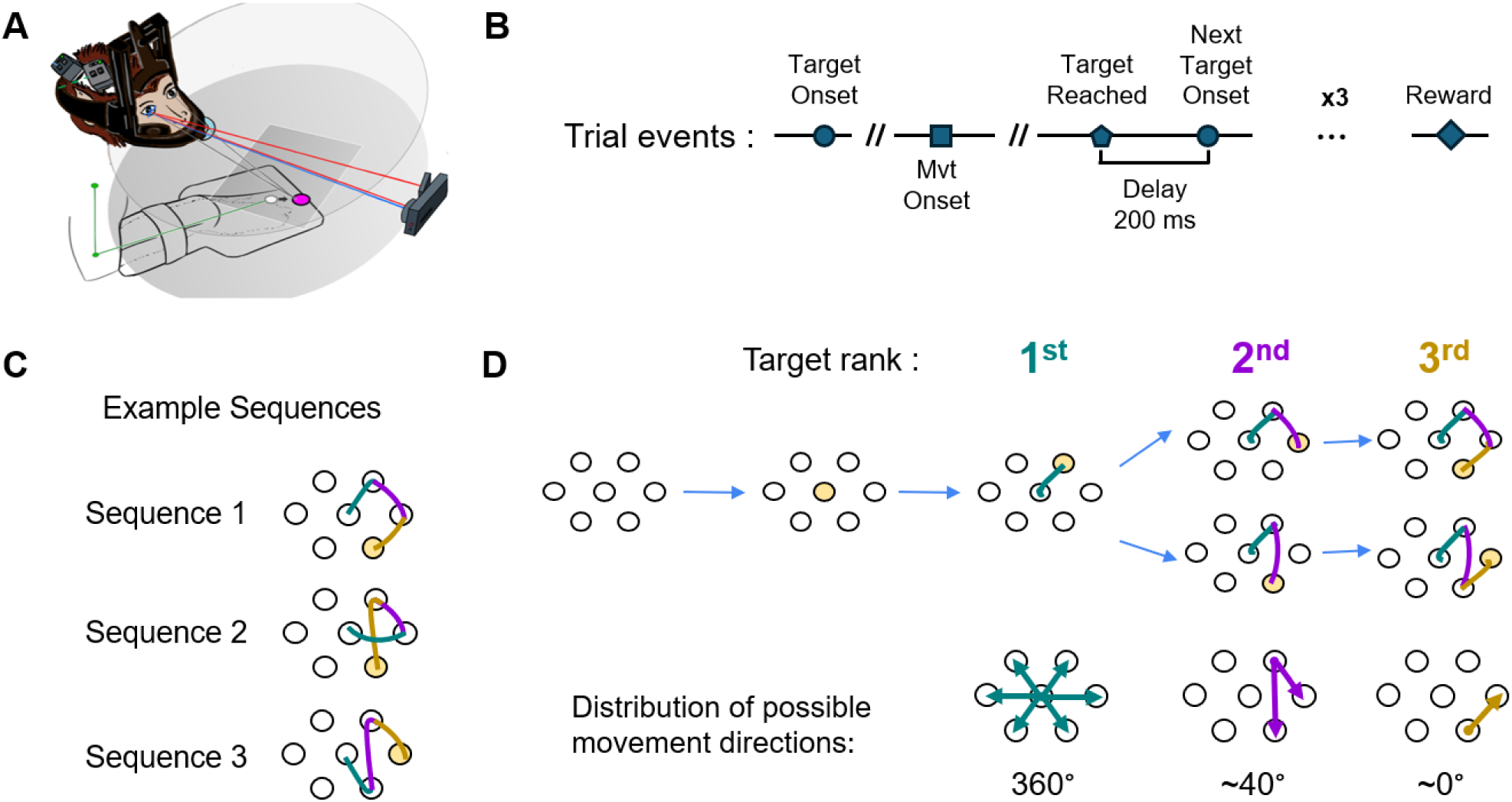
Experimental task design. (A) Two Macaque monkeys were sitting in a primate chair with a mask for head fixation. The monkeys guided a robotic exoskeleton arm towards visual targets that appeared sequentially in a horizontal screen in front of them. (B) Trial design. Trials always started from the initial central target. Once the central target disappeared, a peripheral target appeared simultaneously which indicates the go cue. After a variable reaction time, the monkeys reached the Valid Target Area in which they had to maintain the cursor for 200 ms. Once the reached target disappeared, the next one appeared. Monkeys were rewarded after the completion of three such movements. (C) Three (out of twelve) example sequences that were presented pseudo-randomly on the screen. Each sequence consisted of three reaching movements that formed a complete trial. (D) The predictability of the target location increased within a sequence. The first target could appear in any of the six possible peripheral locations. The second target, given the position of the first, can appear in two possible locations and the third target given the position of the two previous is deterministic.

### Hand and eye movements reveal distinct behavioral strategies

We analyzed 3663 reaching movements from 15 sessions for monkey J and 3249 movements from 14 sessions for monkey E in which both animals had already been extensively trained in the 12 possible target sequences. Despite some inter-trial variability, hand trajectories were consistently oriented towards the target location, even in the initial part of the movement trajectories, indicating that movements were goal-directed (**Figure 2A**, **Figure S.2A**).

**Figure 2.**
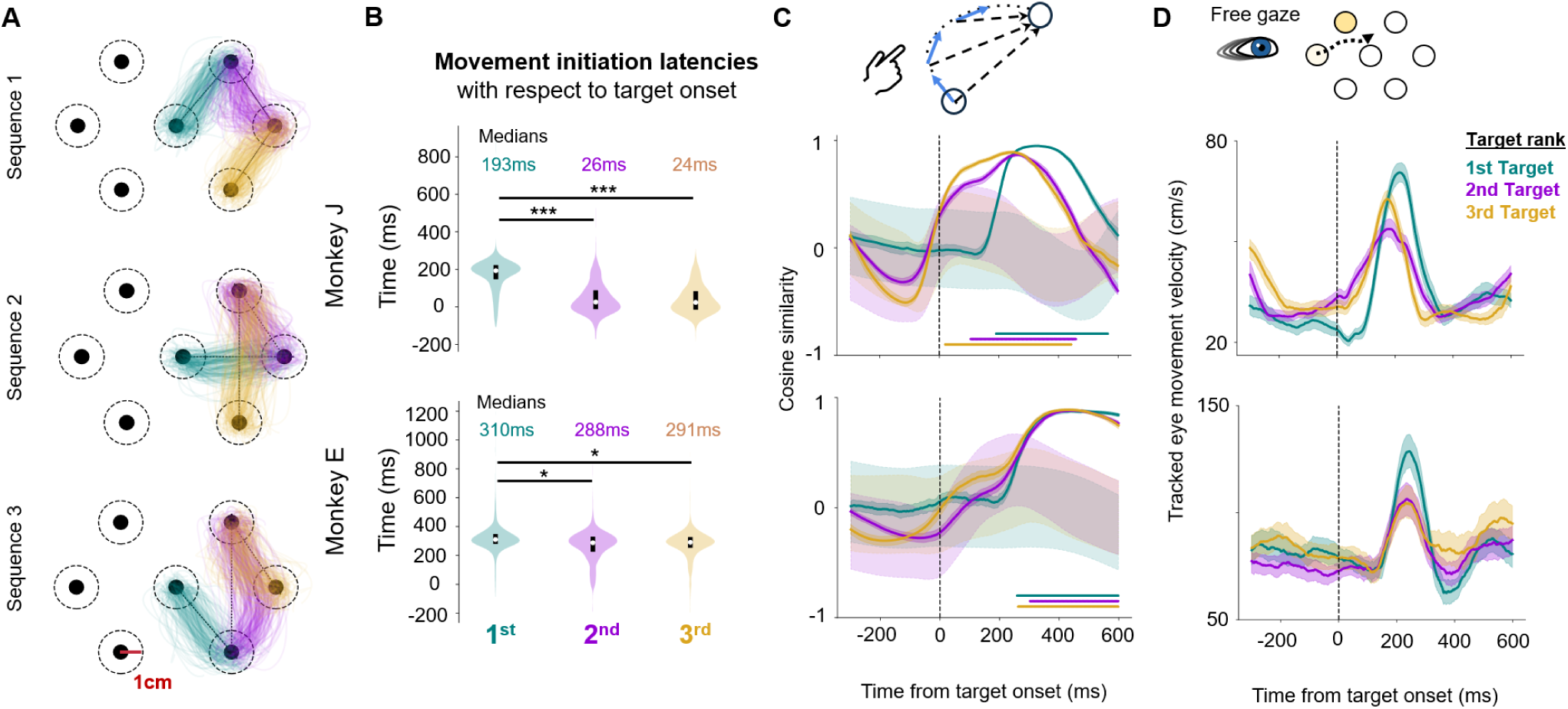
Hand and eye movements reveal distinct visuomotor strategies. (A) Single-trial hand trajectories (Monkey J) color-coded for the 1^st^ (teal), 2^nd^ (violet) and 3^rd^ (gold) movements of the trial for the example sequences of Figure 1B. (B) Movement initiation latencies with respect to the 1^st^, 2^nd^ and 3^rd^ target onset for both monkeys. P values were computed applying Wilcoxon signed rank tests with Bonferroni correction: p < 10^-20^ is noted with *; p < 10^-140^ is noted with ***. (C) Cosine similarity between the instantaneous hand trajectory direction (blue vectors on top schema) and the instantaneous target direction (black dashed vectors on top schema) for each target rank aligned on each target onset. Confidence intervals around the trial-average cosine similarity show the 95^th^ percentile of the estimation of the mean (200 bootstraps). Shaded confidence intervals around zero represent the chance level obtained by computing the similarity after shuffling the target direction and showing the mean ± half standard deviation of the shuffled distribution. Horizontal bars at the bottom mark the significant epochs obtained from a one-sample t-test against chance distribution. (D) Tracked eye movement velocity in the Cartesian coordinate system aligned on each target onset. Confidence intervals show the 95th percentile of the estimation of the mean (200 bootstraps). Wider confidence intervals reflect fewer trials, which resulted from discarding trials with noisy tracking of eye positions.

We first examined whether monkeys anticipated the movement towards a target location as a function of target predictability. Movement initiation latencies with respect to target onset decreased significantly in both monkeys (**Figure 2B**). Significant differences were observed between the 1^st^-2^nd^ and 1^st^-3^rd^ targets. This indicates early movement due to prediction of the timing of the next target appearance, but not necessarily anticipation of the target’s spatial location. Specifically for monkey J, movement initiation toward the 2^nd^ and 3^rd^ targets was simultaneous with, or even earlier than, target onset (**Figure 2B** top, violet and gold distributions). This suggests that monkey J initiated movement preparation before the visual target even appeared, which was possible given the large diameter of the Valid Target Area. Movement execution durations did not vary significantly across targets (**Figure S.2B**), with a slight increase for the 2^nd^ and 3^rd^ targets.

To evaluate whether monkeys anticipated the future target location, we quantified how closely the hand movement direction aligned with the direction of the upcoming target at each moment in time, calculating their cosine similarity (**Figure 2C** top). We observed that cosine similarity for movements toward the unpredictable 1st target increased significantly about 200-300 ms after the target onset (**Figure 2C**), matching each monkey’s movement initiation latencies. The null distribution of trajectory to target alignment by chance was estimated by shuffling target direction (see Methods). The similarity value of movements towards the 2nd and 3rd targets surpassed chance level gradually closer to target onset for monkey J. Interestingly, in contrast to movement initiation latencies, which did not differ between 2^nd^ and 3^rd^ targets, the alignment of the trajectory with the target direction became significant earlier for the 3^rd^ target than for the 2^nd^ (violet and gold horizontal lines in **Figure 2C**), consistent with the greater predictability of the 3^rd^ target. These results indicate that goal-directed movements toward 2^nd^ and 3^rd^ targets were planned in anticipation of these targets’ appearance on the screen. In contrast, cosine similarity values for monkey E increased significantly at very similar time across targets, indicating that the observed decrease in movement initiation latencies was mainly related to learning the timing of task events.

We also examined whether the monkeys’ eye movements showed signs of anticipation. In monkey J, the latency of eye movement onset decreased significantly between 1^st^-2^nd^ and 1^st^-3^rd^ targets (**Figure S.2C**), matching the hand movement initiation latencies. Hand and eye movements occurred at similar latencies after the 1^st^ target, but the hand started moving earlier than the eye at the onset of the 2^nd^ and 3^rd^ targets, as shown by the significant decrease in the single-trial hand-eye onset delays (**Figure S.2D**). Conversely, for monkey E, hand movements occurred after eye movements. In monkey J, the average eye movement velocity increased earlier for predictable than for unpredictable targets (**Figure 2D**). Accordingly, eye movement profiles did not vary across targets for monkey E.

Taken together, both hand and eye movements of monkey J demonstrated strong effects of spatiotemporal movement anticipation when target locations were partially or totally predictable. Conversely, monkey E did not exhibit the same strategy, initiating target-oriented movements at similar latency regardless of target predictability, since movement anticipation was not required to complete the task. This behavioral contrast offers the opportunity to investigate whether corresponding differences are present in the underlying neural activity of the two monkeys.

### Distinct motor neural activity reflects different behavioral strategies

We simultaneously recorded neural activity using multielectrode Utah arrays (Blackrock Microsystems), chronically implanted in the parietal Brodmann area 7A and at the junction of the dorsal premotor and the primary motor cortex (PMd/M1) for both monkeys (**Figure 3A**).

**Figure 3.**
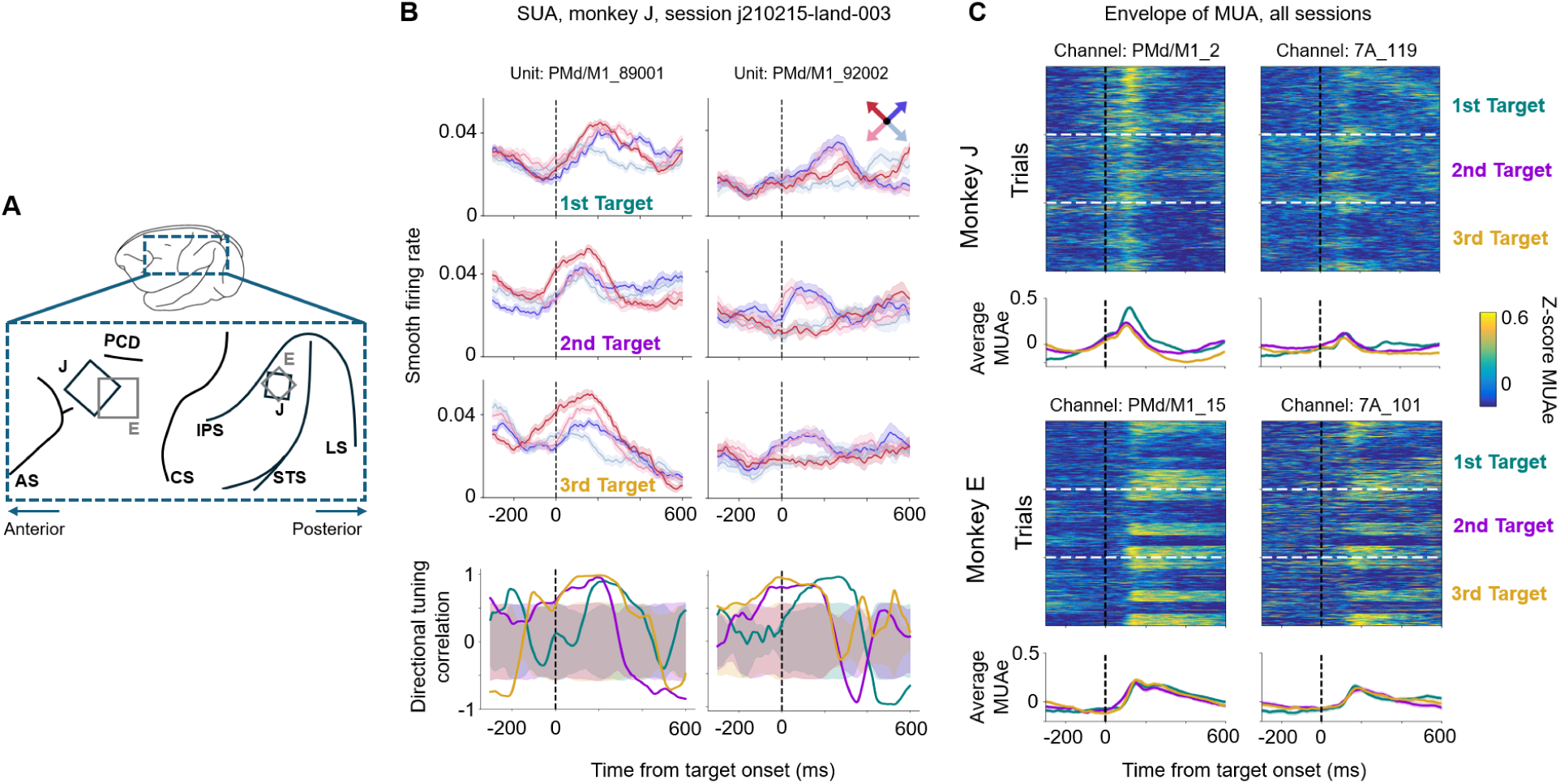
PMd/M1 activity reflects different behavioral strategies. (A) We simultaneously recorded neural population activity from a 96-channel Utah array in PMd/M1 and a 32-channel Utah array in 7A. Schematic representation of the implantation sites of the Utah arrays for the two monkeys in a common reference along the anterio-posterior axis. J,E - the monkeys’ initials, PCD-Precentral dimple, AS-Arcuate sulcus, CS-Central sulcus, IPS-Intraparietal sulcus, STS-Superior Temporal Sulcus, LS-Lunate Sulcus. From the high-pass filtered extracellular voltage of each session, we sorted spikes into single units and computed the envelope of the MUA for each channel of each array. (B) Single-unit activity (SUA) from two example units from an example session. Top. Firing rates convolved with a Gaussian kernel averaged in each of the four movement directions that are repeated across target ranks. Each row represents the activity aligned on the 1^st^, 2^nd^ and 3^rd^ target rank. Each column shows the results for one unit. Bottom. Correlation of directional tuning at each time point with the mean directional tuning in the window 100-300 ms after 1st target onset. Confidence intervals represent the chance level obtained by computing the correlation after shuffling the directional tuning after the 1^st^ target onset. (C) Z-scored MUA in single trials concatenated across all sessions aligned on target onset for one example channel in each area for both monkeys. Trials are grouped in the three target ranks (the three groups are separated by dashed white lines). Within each target rank, trials are sorted based on movement direction. Line plots below the heatmaps show the trial-average MUA in the three target ranks.

To examine whether neural activity encoded movement direction earlier for predictable than unpredictable targets, we first evaluated whether directional tuning in single-unit activity (SUA) of PMd/M1 emerged earlier when targets could be predicted. When targets were unpredictable (1^st^ target), directional tuning, reflected by higher firing rates for each unit’s preferred directions, emerged about 200 ms after target onset in both monkeys. But when targets were predictable (2^nd^, 3^rd^ targets), directional tuning emerged even before target appearance (two example neurons are shown in **Figure 3B**) exclusively in monkey J who exhibited anticipatory behavior. To quantify this temporal shift, we defined the unit’s “template” tuning vector as the average firing rate in the interval 100–300 ms after the 1^st^ target onset, and we computed the correlation between this vector and the instantaneous firing rate across directions at each timepoint. We found that the correlation with the template tuning vector became significant before the onset of predictable targets (violet and gold lines in bottom plots of **Figure 3B**).

We next examined whether anticipatory representations observed in single units were reflected in population-level activity. We calculated the envelope of the Multiunit Activity (MUA, Buchwald et al., 1965), which provides a mesoscopic measure representing the integrated firing activity of the local neuronal population around each electrode. Such population-level signal is particularly suited for the study of interareal mesoscopic interactions, because MUA preserves substantial movement-related information and has been successfully used for movement decoding (Stark & Abeles, 2007) and interareal interactions (Celotto et al., 2023). Moreover, its greater reproducibility across recording days allows the concatenation of trials across sessions, thereby increasing the signal-to-noise ratio and number of trials available per direction.

For monkey J, MUA amplitude in PMd/M1 decreased significantly across single-trials when targets were predictable (one example channel in **Figure 3C**, statistics across channels in **Figure S.3A**). This difference in the amplitude of MUA peaks across targets could not be explained by the starting position from which the monkeys initiated their hand movement (**Figure S.4**), which also changed during the sequence. Conversely, MUA modulation in PMd/M1 for monkey E did not show similar differences across targets, indicating that the effect couldn’t be attributed to simpler effects, such as elapsed time in the sequence or reward-related modulations. MUA in parietal area 7A didn’t exhibit significant amplitude differences across targets in both animals (**Figure 3C**, **Figure S.3A**). The latency of the MUA peak shifted slightly earlier in both areas of both monkeys, potentially due to temporal prediction of the ‘go’ (**Figure S.3B**).

Our findings show that, when targets were predictable, directional tuning in single units of PMd/M1 emerged earlier and the amplitude of mesoscopic neural population activity decreased significantly for monkey J, who exhibited behavioral anticipation (**Figure 2**). In contrast, neural activity for monkey E, who waited for the visual cues before moving, did not show differentiated modulations across targets. Together, these results reveal two different neural activity profiles that describe the two corresponding behavioral strategies for solving the task.

### Preparatory motor and parietal dynamics track behavioral strategy

To assess whether neural population activity tracked the behavioral strategy, we examined whether the emergence of neural representations of the planned direction shifted before the visual target onset during motor preparation for predictable targets. In the dynamical systems perspective (Shenoy et al., 2013), preparatory motor activity is thought to set the initial conditions of a dynamical system that evolves to generate desired movements. Within this framework, anticipation ought to bring preparatory dynamics to these initial conditions before sensory information arrives, allowing the system to lead efficiently to appropriate anticipatory movement trajectories. In particular, we hypothesized that prior information about the movement, both in motor and parietal responses, would lead to a faster reaching of the initial conditions, even before the visual stimulus appears. By contrast, in absence of motor planning prior to the visual stimulus, movement trajectories should occur strictly after sensory information arrives.

To do so, we represented multichannel activity at a given time as a point in a high-dimensional space, where each dimension corresponded to a single channel. We performed a Linear Discriminant Analysis (LDA, widely used in electrophysiological studies, Meyers et al., 2012; Boulay et al., 2016; Yiling et al., 2023; Wang et al., 2024) to identify two discriminant axes that best separated the high-dimensional MUA based on the instructed movement direction (**Figure 4A**). To specifically extract the motor planning component of the neural activity, we trained the model to discriminate single-trial MUA based on direction during the first 200 ms following the 1st target onset. To assess whether the axes describing the neural activity around the unpredictable target provided a suitable discriminative space for the direction of predictable targets, we projected MUA from the 2nd and 3rd targets onto these axes. We observed that shortly after the 1^st^ target onset, PMd/M1 projections on DA1 split left from right movement directions (discrimination of red from blue palette colors respectively, **Figure 4B**) and projections on DA2 split top from bottom directions (discrimination of dark from light colors respectively, **Figure 4B**), as expected from training the model on that epoch. Such representation emerged prior to the onset of the following targets in the sequence for monkey J (**Figure 4B** middle and right columns), whereas for monkey E it appeared at similar latencies across target ranks, after target onset (**Figure S.5A**). Since leftward and rightward horizontal movements did not occur for the 2nd and 3rd targets, only four repeated directions are shown in **Figure 4B** (middle and right columns). Upward and downward vertical movements, which were absent from the training data, were projected between their neighboring directions (top-right/top-left and bottom-right/bottom-left, respectively) in both monkeys (grey and black projections in **Figure S.5A**). Similarly, neural activity in 7A was clustered in the same neural space locations for the same movement directions across target ranks (**Figure S.5B**).

**Figure 4.**
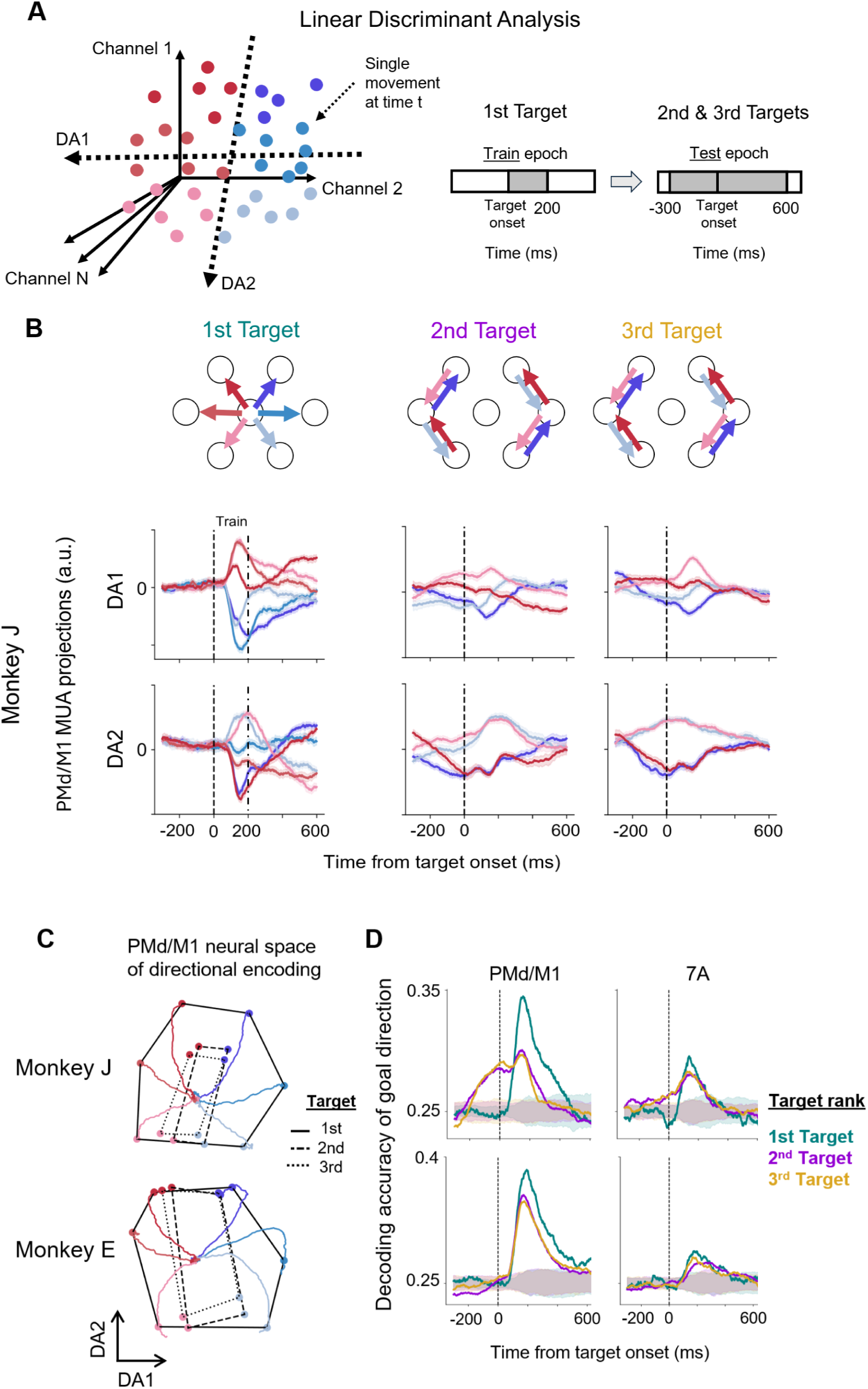
Preparatory motor and parietal dynamics track behavioral strategy. (A) Linear Discriminant Analysis (LDA). We applied LDA in each area to get two discriminant axes that maximally separate the trajectories in the six different movement directions of the center-out movements toward the 1^st^ target. We trained the LDA and got the new dimensions based on the first 200ms after the 1^st^ target onset (unpredictable upcoming movement). We then projected the MUA of the 2^nd^ and 3^rd^ segment onto the same axes aligned on each target onset. (B) PMd/M1 MUA projections (Monkey J). Each of the two rows shows one discriminant axis (DA1 and DA2), while each of the three columns indicates the epoch around the 1^st^, 2^nd^ and 3^rd^ target respectively. Each line shows the average projection across movements to the same direction (same color code as in A) in arbitrary units. Confidence intervals show the 95th percentile of the estimation of the mean (200 bootstraps). (C) The PMd/M1 neural space of directional encoding for both monkeys. Colored lines show the evolution of the neural trajectory in the space defined by DA1 and DA2 after the 1^st^ target onset (same color code as in A). The colored dots (one per direction) show the projection at the moment of the maximum variance across directions for each target rank. Black lines connect these dots to dissociate projections belonging to the same target rank (1^st^ target - solid line; 2^nd^ target dashed line; 3^rd^ target dotted line). (D) Decoding accuracy of LDA predicting the movement direction for each of the three epochs, around the 1^st^ (teal), 2^nd^ (violet) and 3^rd^ (gold) targets. To cross-validate the accuracy of the 1^st^ target, we trained the model in 80% of the segments and computed the accuracy on the remaining 20%. The shaded intervals show the distribution of accuracy by chance obtained by LDA models trained on shuffled direction labels. Chance was around 0.25 since the decoder was trained to classify the four possible directions that can be repeated in all target ranks (i.e. top-right, top-left, bottom-right, bottom-left).

To evaluate the geometry of the projections, we depicted the neural representations as a hexagonal space created by the projections on DA1 and DA2 at the moment of the maximum variance across directions (**Figure 4C, Figure S.5**). The 2D neural trajectories converged to the same state each time monkeys planned a movement towards a specific direction in all target ranks. This confirms that neural representations of directional tuning in PMd/M1 during motor planning were preserved in the same latent space irrespective of target predictability and hand position at movement initiation.

Importantly, the divergence of the neural trajectories, marking the initiation of motor planning (Churchland et al., 2006), occurred prior to the onset of the 2^nd^ and 3^rd^ predictable targets (**Figure 4B** middle and right columns). To quantify the re-emergence of these neural representations, we computed the cross-validated accuracy of the LDA predicted movement direction. Based on the MUA directional modulations, the LDA classifier predicted the movement directions that were common across target ranks (i.e. top-right, top-left, bottom-right, bottom-left). Chance level accuracy distributions were calculated from the predictions of models trained on shuffled direction labels, separately for each target (see Methods).

In monkey J, the prediction accuracy became significant about 50 ms after the 1^st^ target onset (unpredictable) in both areas (**Figure 4D**). Interestingly, we decoded forthcoming movement direction 200 ms prior to the presentation of the 2^nd^ and 3^rd^ targets (predictable) in PMd/M1 (and 50 ms prior in 7A). However, for movements toward predictable 2^nd^ and 3^rd^ targets in which monkey J did not exhibit motor anticipation (reaction time > 150 ms in **Figure 2B**), decoding accuracy became significant only after target onset, as for the 1st target (**Figure S.6**). Thus, the temporal dynamics of directional neural representations reflected the behavioral strategy employed, even within the same monkey.

Decoding performance for the 2^nd^ and 3^rd^ targets (test set) was lower than for the 1st target (train set), consistent with the neural space projections, where the quadrilaterals of 2^nd^ and 3^rd^ targets exhibited less separation across directions than the hexagon corresponding to the 1^st^ target (**Figure 4C**). This reduction was also consistent with the MUA amplitude decrease in monkey J (**Figure 3C**). Accuracy in 7A was lower than PMd/M1, but remained significantly above chance, since activity corresponding to the same movement direction clustered at similar locations in the neural space across target ranks (**Figure S.5B**).

In monkey E, decoding accuracy emerged at similar latencies after target onset in both areas, irrespective of target rank. Decoding accuracies were comparable across target ranks, consistent with the similar separation of the neural trajectories (**Figure 4C**) and comparable MUA amplitudes (**Figure 3C**).

These findings demonstrate that when movements were anticipated, preparatory dynamics of directional tuning emerged before the presentation of the visual cues, indicating that motor planning was initiated by internal predictions rather than external stimuli. Movement anticipation shifted the emergence of the directional tuning earlier not only in PMd/M1, but, interestingly, also in area 7A (**Figure 4D**; **Figure S.5B**), indicating earlier planning of, not only hand, but also eye movements in anticipation of predictable visual targets. By contrast, in absence of motor anticipation, preparatory dynamics occurred only after visual target onset, at similar latency regardless of target predictability. These results indicate that preparatory dynamics in both motor and parietal areas flexibly reflect the visuomotor strategy employed.

### Bidirectional, rather than strictly serial, parieto-motor interactions are preserved across strategies

Building on the emergence of individual motor and parietal preparatory dynamics, we investigated whether their statistical interactions were uni- or bidirectional. Based on traditional models of hierarchical sensorimotor processing (Scott, 2012), encoding of direction –representing planning– in parietal regions is expected to precede motor dynamics, exhibiting thus unidirectional feedforward parieto-motor interactions. Alternatively, based on recent evidence of brain-wide distributed processing (Steinmetz et al., 2019), interactions are expected to be bidirectional: feedforward parieto-motor transformations and feedback efference copies of the motor command to parietal areas, potentially used for internal modeling (Wolpert et al., 1995; Todorov & Jordan, 2002).

To characterize these interactions, we employed a novel information-theoretic measure, the Feature-specific Information Transfer (FIT, Celotto et al., 2023) that has already been applied to electrophysiological recordings to quantify task-related directional interactions between neural signals (Lemke et al., 2024; Combrisson et al., 2025). The main advantage of FIT, over alternatives such as Granger Causality (Granger, 1969), lies in its ability to quantify directed interdependencies between time series, while explicitly linking the conveyed information to a task-relevant variable. We computed FIT about the instructed movement direction between the MUA of channel pairs from the two areas. FIT increased when information about the forthcoming direction in MUA from a source channel was shared with that of a target channel within delays of up to 100 ms (**Figure 5A**). Importantly, FIT captured the additional information about movement direction that was present in the past of the source and not in the past of the target activity. Using Canonical Correlation Analysis, we verified that the correlated activity between past and present MUA from channels of the two areas reflected aligned representations of movement direction (**Figure S.7**), as indicated by the consistent ordering of the four directions in **Figure 5A**.

**Figure 5.**
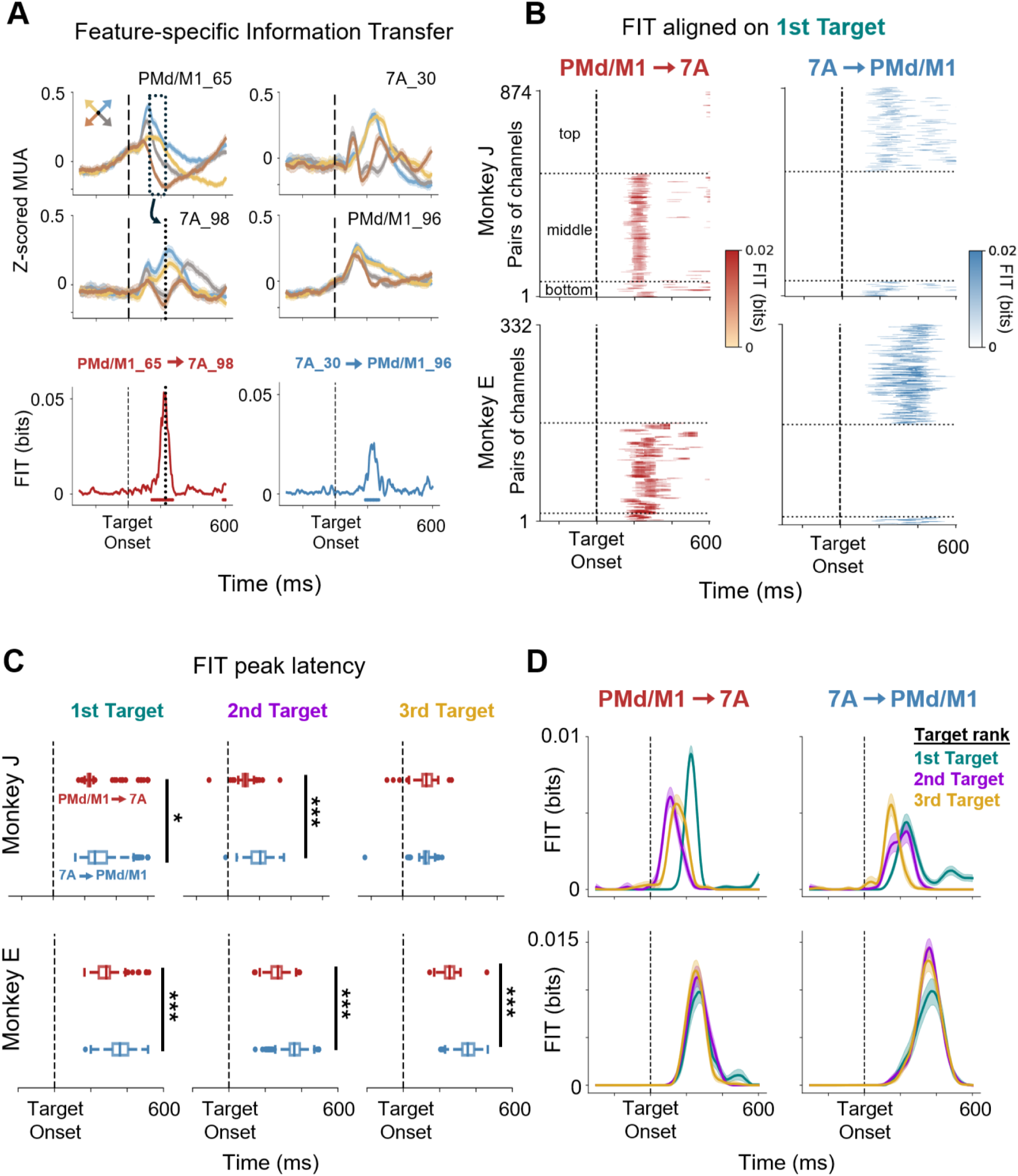
Bidirectional motor-parietal interactions are preserved across strategies. (A) Top. The MUA across the four directions that are common across target ranks for two example channel pairs. Bottom. The Feature-specific Information Transfer (FIT) about the movement direction for each channel pair. In the first pair (PMd/M1_65, 7A_98), the split across directions in PMd/M1 preceded the split in 7A, thus the FIT increased in the direction from PMd/M1 to 7A. In the second pair (7A_30, PMd/M1_96), conversely, the FIT increased in the direction from 7A to PMd/M1. (B) The FIT for all significant channel pairs around the 1^st^ target onset. Each row of each subplot shows the FIT (in bits) of one pair of channels between the two arrays. Clusters were considered significant if p-values were smaller than 0.05 for more than 50 ms consecutively; only significant FIT clusters are shown. Pairs of channels are arranged in three groups depending on whether they have significant FIT in both directions (lower group), only in the direction PMd/M1 → 7A (middle group) or only in the direction 7A → PMd/M1 (upper group). (C) Distribution of latencies of the FIT peak for all significant pairs of channels in each direction for each target. Solid black lines indicate the significant differences between the distribution medians using permutation testing. Median difference smaller than 50ms is noted with *, while larger than 50 ms is noted with ***. Dots outside boxplots show the channel pairs whose latency lies outside of the distribution. (D) The FIT averaged across significant channel pairs in the direction PMd/M1 → 7A and in the direction 7A → PMd/M1 across targets. Confidence intervals show the 95th percentile of the estimation of the mean (200 bootstraps).

Out of all channel pairs between the PMd/M1 and 7A arrays for which both channels showed significant directional modulation, 874 pairs (32.1%) exhibited significant FIT around the 1st target onset in monkey J, compared with 332 pairs (9.8%) in monkey E. Significant movement-direction FIT values rarely formed clusters in localized anatomical positions; they were rather distributed across the array with no clear spatial gradient or pattern (**Figure S.8**). Percentages of channel pairs with significant FIT around the 2nd and 3rd target onset were similar (**Figure S.9**). Only a limited number of channel pairs showed significant FIT in both directions (bottom group in **Figure 5B** & **Figure S.9**). Significant FIT in a single direction, either PMd/M1→7A (middle group) or 7A→PMd/M1 (top group), was more common. We observed that the number of significant channel pairs was quite balanced across directions for both monkeys (**Figure 5B**): in monkey J, 40.4% were 7A→PMd/M1 and 52.4% were PMd/M1→7A; in monkey E, 50.3% were 7A→PMd/M1 and 45.8% were PMd/M1→7A. These results do not support a dominant feedforward parieto-motor organization, which would predict a clear predominance of 7A→PMd/M1 interactions.

The temporal lead/lag relationships between encoding of direction in motor and parietal responses occurred at variable latencies across significant channel pairs. Median latency differences (computed as PMd/M1→7A minus 7A→PMd/M1) indicated that motor-parietal interactions preceded parieto-motor interactions (**Figure 5C**). For monkey J, the median differences were 35 ms (1st target), 95 ms (2nd target), and 5 ms (3rd target); for monkey E, they were 75 ms, 90 ms, and 100 ms, respectively. These findings indicate that the encoding of forthcoming movement direction is not organized as a strictly unidirectional parietal-to-motor process, but instead involves reciprocal motor–parietal interactions, with directional information emerging earlier in motor than in parietal responses.

Finally, we examined whether increased predictability temporally advanced not only neural responses within parietal and motor areas (**Figure 4D**), but also the timing of their directed interactions. Specifically, we tested whether the directed interactions were preserved but occurred earlier in time. This would indicate that anticipation modulates interareal network dynamics, not just the onset of activity within individual regions.

Indeed, for monkey J who behaviorally anticipated the movements, the average FIT peak occurred earlier when the monkey moved to predictable targets in contrast to movements to unpredictable targets (**Figure 5D**, **Figure S.10B**). The latency shift indicates that the co-activations were related to the anticipatory hand/eye movements, rather than to the visual stimulus, which would require time-locked responses. Also, the FIT amplitude decreased consistently across channel pairs, not resulting from averaging misaligned information values due to earlier movements (**Figure S.10A**). Furthermore, the temporal lag between interareal co-activations remained constant across targets of different predictability (**Figure S.10C**, PMd/M1→7A). This could suggest that, although the system adjusts the timing of directional encoding in each individual area to plan movements in anticipation of visual stimuli, the relative timing between motor and parietal responses is preserved. In monkey E, who did not show behavioral anticipation, the effects did not differ systematically across targets.

Overall, these findings demonstrated that statistical interdependencies between motor and parietal dynamics were bidirectional, and not strictly serial, and flexibly reflected behavioral strategies by temporally advancing during motor anticipation, aligning with a reciprocal model of distributed information processing that extends beyond classical models of unidirectional sensorimotor transformation.

## Discussion

In this study, we investigated the properties of motor-parietal interactions during motor planning and their flexibility in adapting to various environmental constraints. We first showed that the neural representations underlying motor planning, reflected by directional tuning of single neurons and MUA, emerged earlier for anticipated movements, preceding the appearance of the visual target. We then showed that movement-direction-specific information transfer across PMd/M1 and 7A was bidirectional with no serial parietal followed by motor activations. Overall, our findings indicate that motor–parietal interactions during motor planning of different visuomotor strategies are not organized in a purely feedforward direction, but instead reveal reciprocal interactions. Moreover, this pattern of interactions was preserved during motor anticipation. Within this framework, PMd/M1 and area 7A represent crucial nodes of a distributed fronto-parietal network that reshapes the timing of preparatory activity to support flexible motor planning during sequential reaching.

### Flexible motor planning and anticipation

It is well established that neurons in PMd and M1 are activated before movement initiation, to prepare the upcoming response to task-related visual stimuli (Confais et al., 2012; Crammond & Kalaska, 2000; Riehle & Requin, 1989). When the upcoming target location is partially or fully predictable, decoding accuracy of the movement direction from PMd/M1 during the preparatory delay between target onset and go signal improves significantly (Bastian et al., 2003; Rickert et al., 2009; Rostami et al., 2024). In our task, one monkey anticipated and one did not, therefore allowing us to test if there were correlates of this behavioral contrast in the motor and parietal neural activity as well as their interactions. We found that in the monkey showing movement anticipation, the MUA in PMd/M1 decreased significantly, from the unpredictable to the predictable targets (**Figure 3C**, **Figure S.3A**). This is in agreement with single-neuron spiking activity in PMd/M1 before movement onset. When movement direction is known in advance, neuronal activity is lower than when no prior information about movement direction was provided (Riehle & Requin, 1989). In both partially and fully predictable targets (2nd and 3rd), monkeys could broadly anticipate the movement direction and correct the trajectory once the visual target appeared, as reflected in the cosine similarity of the 2nd target, which increased more slowly than for the 3rd target (**Figure 2C**). The observed MUA amplitude decrease may reveal an interesting aspect of our results that can be related to one expectation of the predictive coding theory (Rao & Ballard, 1999). According to this hypothesis, such amplitude decrease for predictable targets is consistent with the proposed attenuation of neural activations due to lateral inhibition when visual stimuli are predictable (Rao and Ballard, 1999), that we extend here to the premotor cortex in which visual target information is also represented (Cisek and Kalaska, 2002). However, we may not exclude alternative explanations, such as distinct neural encoding of center-out vs peripheral reaches during sequential movements.

In the dynamical systems perspective, PMd/M1 activity during movement preparation does not simply reflect the future movement features. Instead, it needs to reach an optimal state that determines the initial conditions from which the neural dynamics will evolve to produce the desired movement (Churchland et al., 2006, 2010, 2012; Churchland & Shenoy, 2024; Kaufman et al., 2014; Shenoy et al., 2013). Consistent with these research lines, we showed that the initial conditions defined by the preparatory activity in PMd/M1 remained the same for movements in the same direction, irrespective of the visual target predictability (dots that form the spaces in **Figure 4C**). However, when the future target location was anticipated, the motor planning initiation, here defined as the neural trajectories’ split, was shifted before the visual target’s appearance (**Figure 4B**,**D**). This led to earlier and correct movement execution with minimal reaction time for predictable targets (**Figure 2B**), providing a signature of anticipation at the neuronal level.

In this task, the visual target onset also served as the go signal with no delay separating these two events. However, the consistent accuracy of hand trajectories, even at their initial stages (**Figure 2A**, **Figure S.2A**), indicates that brief motor preparation still occurred between the sequential movements-even the ones with the shortest reaction time. This process may have involved either planning during the dwelling phase at each target or simultaneous preparation at the final stages of the previous movement (Ames et al., 2019). These results align with previous findings showing that preparatory events are conserved irrespective of the presence or absence of a delay period and even during reactive movement corrections, where reaction times are typically very short (Ames et al., 2014, 2019; Lara et al., 2018).

### Involvement of area 7A in motor planning

Areas of the posterior parietal cortex (PPC) have been linked to forming movement intentions (Andersen & Buneo, 2002), participate in motor planning (Cui & Andersen, 2007, 2011), and encode goal and movement angle (Mulliken et al., 2008b), serving as a critical node in the forward internal model for sensorimotor control (Todorov & Jordan, 2002; Wolpert et al., 1995). Although area 7A (or area PG in the Von Economo nomenclature) was traditionally viewed as an associative visuospatial area (Hyvärinen & Poranen, 1974; MacKay, 1992; Rolls et al., 2022), recent evidence has revealed that neurons in area 7A encode movement direction towards static (Heider et al., 2010) and moving targets during interception (Li et al., 2022). Critically, several reports have emphasized the functional relevance of area 7A in reaching during complex tasks that require continuous eye-hand coordination and visuospatial analysis (Battaglia-Mayer & Caminiti, 2019), such as the sequential reaching task of increased predictability studied here. In a task that required eye-limb coordination during locomotion, area 7 neurons in cats encoded the relative distance to obstacle (Marigold & Drew, 2017), indicating that this area is involved in computations related to upcoming movements. In our task, we show that, when the upcoming target was anticipated, low-dimensional population dynamics in area 7A exhibited anticipatory directional tuning. These dynamics enabled the decoding of a predictable target location even before its appearance on the screen (**Figure 4D**), extending previous works showing that PPC ensembles can encode the reach endpoint (Musallam et al., 2004) and even complete hand trajectories (Mulliken et al., 2008a). Although significant, the decoding accuracy was modest in comparison to other studies (Ganguly & Carmena, 2009; Shanechi et al., 2012). This difference may arise from a couple of factors acting in combination. First, unlike these studies, our decoder was trained on MUA, which is likely noisier than SUA for single-trial decoding. Second, the sequential task used in our study is more complex than standard center-out reaching tasks and likely introduces some variability due to differences in initial position for movements in the same direction. As a result, we consider the significance of the decoder, which represents the moments at which the neural trajectories maximally split, as the key factor for interpreting the results.

Mechanistically, such neural selectivity in 7A could be driven by a saccade to the position of the target (Andersen et al., 1987, 1990; Barash et al., 1991), prior to its presentation. Consistent with this possibility, decoding accuracy of the upcoming direction in 7A (**Figure 4D**, **Figure S.5B**) temporally coincided with the eye velocity profiles (**Figure 2D**), suggesting that the low-dimensional dynamics in 7A may partly reflect the direction of the anticipatory saccade. Establishing a direct relationship between 7A activity and eye behavior was challenging, because the task was performed under free-gaze conditions. In a paradigm that would require fixation to the center followed by a saccade to each target, we would predict that 7A neural representations would allow stronger future direction decoding than observed here (**Figure 4D**). Nevertheless, we would not expect this manipulation to alter the reciprocal pattern of motor-parietal interactions (**Figure 5**). Our finding that motor-parietal preceded parieto-motor interactions argues against a purely oculomotor account in which 7A passively reflects eye movement execution.

An alternative explanation is that 7A may have received movement-related information about the direction in the form of an efference copy of the motor command (Todorov & Jordan, 2002; Wolpert & Flanagan, 2001). In fact, previous studies have shown that neural representations in area 7A encode information about the forthcoming movement, rather than visual stimuli, indicating a strong link with motor planning (Li et al., 2022). Consistent with this evidence, we found that interactions from PMd/M1 to 7A preceded interactions from 7A to PMd/M1 (**Figure 5C**). Therefore, these results indicate that area 7A could be recipient of efference motor copies and function as an active node within a reciprocal fronto-parietal network that enables predictive motor preparation, rather than as a relay along bottom-up pathways for visuomotor transformations.

### Functional roles of motor–parietal interactions

Classical models of parieto-frontal circuits typically emphasize the bottom-up sensorimotor transformations of visual stimuli from retina-to limb-centered coordinates to enable the motor system to issue motor commands and guide the appropriate muscles (Goodale & Milner, 1992; Wise et al., 1997; Rizzolatti et al., 1997; Snyder et al., 1998; Caminiti et al., 1998; McGuire & Sabes, 2009; Rizzolatti et al., 2014). Area 7A inputs to motor regions have been linked to such coordinate transformations not only to initiate visually guided movements (Buneo & Andersen, 2006; Scott, 2004), but also to estimate their sensory consequences (Blakemore & Sirigu, 2003) and support online corrections (Archambault et al., 2015; Sarlegna & Mutha, 2015). Interestingly, even imagined motor goals, hand trajectories and movement types could be decoded from PPC neurons (Aflalo et al., 2015).

Despite the suggested involvement of area 7A in motor processes (Heider et al., 2010; Li et al., 2022) and the reciprocal anatomical connections with premotor areas (Markov et al., 2014), the functional relevance of motor-parietal interactions during cognitive motor behavior remains relatively unclear. Thus far, influential studies have investigated fronto-parietal interactions during attentional processes (Buschman & Miller, 2007), decision-making (Pesaran et al., 2008) and working memory (Salazar et al., 2012). Prefrontal inputs specifically in area 7A appear to support high-level executive functions (Crowe et al., 2013; Merchant et al., 2011), whereas premotor inputs to the parietal reach region have been linked to goal retrieval from working memory (Martínez-Vázquez & Gail, 2018). Here, we examined whether and how motor-parietal interactions were modulated during different visuomotor behaviors that required continuous eye-hand coordination. The observed influence of anticipation on the motor-parietal network (**Figure 5D**) is in line with previous findings suggesting that PPC is involved in anticipatory spatial attention of visual stimuli (Bressler et al., 2008), anticipatory motor planning (Snyder et al., 2006) and motor control (Krause et al., 2014).

Recent theories postulate that the neocortex operates under an active predictive coding framework (Rao, 2024). In this model, cortical regions across the hierarchy estimate sensory states and motor actions while anticipating their consequences, with higher-order areas providing feedback that modulates the dynamics of lower-order areas. For instance, neurons in 7A encode latent sensory states, such as the hand displacement from goal (Lakshminarasimhan et al., 2023), while rotational population dynamics in motor areas drive muscle activity to generate movements, evolving from the initial conditions set by the preparatory state (Churchland et al., 2012; Churchland & Shenoy, 2024). Lines of empirical evidence, supporting this theory, have shown that flexible sensorimotor decisions rely on the dynamic interplay of feedforward and feedback information flow within the fronto-parietal network (Pesaran et al., 2008; Siegel et al., 2015). Consistent with these reports, we found bidirectional, rather than serial, statistical interactions between PMd/M1 and 7A during planning of reaching movements. In line with our findings, previous studies have reported that task-related representations emerge simultaneously in primary motor cortex and area 7A during maze solving (Crowe et al., 2004b). Additionally, we have shown that these interactions were preserved during anticipation (**Figure 5D**, **Figure S.9**) and exhibited a corresponding temporal advance, following the shift of the local neural activity. In other words, although activity in both areas shifted earlier in time (**Figure 4D**), the relative interareal delay was preserved, enabling the detection of significant FIT about movement direction for movements to predictable targets. Irregular temporal shifts between the two regions would, otherwise, disrupt this lead–lag structure and result in non–significant estimates (Celotto et al, 2023). Together, these results provide further evidence that PMd/M1 and area 7A participate in a common functional network supporting eye–hand coordination during reaching movements. During motor anticipation, motor-parietal statistical interactions are preserved and temporally advanced, providing further evidence supporting their functional link.

To conclude, our study revealed reciprocal and temporally structured fronto-parietal interactions, aligning with a ‘heterarchical’ model of distributed information processing that extends beyond the traditional hierarchical models of feedforward sensorimotor transformation. These findings provide a principled framework for further research with causal experimental manipulations to elucidate whether disruption of PMd/M1–7A interactions can deteriorate eye-hand coordination during flexible motor performance.

## Materials and Methods

### Experimental setup

During training and recordings, two rhesus monkeys (Macaca Mulatta, one male, Monkey J, one female, Monkey E) were sitting in a Kinarm primate chair (BKin technologies), where they had to move their right arm guiding an articulated exoskeleton in the horizontal 2D plane. A homemade face mask was used to fixate the monkeys’ head and channel the water reward via a tube. The exoskeleton recorded the hand positions, along with elbow and shoulder positions, while the eye positions were recorded by the Eyelink 1000 (SR research), both at 1kHz sampling rate. A horizontal monitor was placed in front of the monkey to display the task (**Figure 1A**). The monkey could move the exoskeleton that supported its arm below the monitor and the cursor that represented the instantaneous position of the monkeys’ hand was appearing in the monitor as a white dot.

### Behavioral task

The monkeys were trained to perform a visually-guided sequential reaching task consisting of three successive hand movements (**Figure 1B,C**). A visual target (0.325 cm radius) appeared in the center of the screen to mark the beginning of the trial. The monkey guided the hand feedback cursor (0.3 cm radius) into the central target and held it there for 150ms for the target to disappear from the screen. At the same time, a target appeared in one of the six peripheral positions that form a hexagon around the central target. These two simultaneous events served as a ‘go’ signal instructing the monkey to release its current position, to move the cursor towards the active visual target (center-out movement) and to remain with a Valid Target Area for 200 ms (dwelling time after reaching each target). The Valid Target Area was defined as an invisible circle with a logical radius of 1 cm around the target and established to improve the monkeys’ motivation to anticipate the next movement. Thus, during the dwelling time, the monkeys were allowed to move within this circle of 1 cm radius around the target. This allowed the monkeys to initiate their hand movement in preparation of the next reach while staying within the Valid Target Area. When that target disappeared, the next peripheral target appeared (peripheral movement). Once monkeys completed a sequence of three movements (one center-out and two peripheral ones), they received the water reward that marked the end of the trial. If the monkeys did not reach the active target in less than 1000 ms, or if they moved the cursor outside of the Valid Target Area without waiting for the previous target to disappear, the trial was aborted. In this case they had to restart the same sequence, to avoid biases due to the monkeys’ preferences on specific sequences. In total, there were 12 possible sequences of targets that were presented pseudo-randomly (**Figure S.1**). One complete session consisted of 120 trials (10 trials for each sequence), depending on the motivation of the animal. In this study we used data from 15 sessions for monkey J and 14 sessions for monkey E, spanning a total period of about three consecutive weeks for each monkey.

Critically, the 2 monkeys were overtrained to perform the task with the same 12 sequences and could exploit the fact that the predictability of the next target location increased within the sequence (**Figure 1D**). Namely, after the monkey fixated in the central target to initiate the trial sequence, a target would appear in one out of the six targets that formed the hexagon around the central target (6 possible positions distributed over a range of 360°). Then, given the first target position, the next target could appear in only one out of two positions, and, finally, given the previous two target positions, the final target could appear in a single position. In this study, we excluded trials that followed an error trial (due to the same sequence being repeated), in order to ensure that the first movement (center-out) was always unpredictable. The percentage of trials that were accomplished in the first attempt (without errors) was 74.4% for monkey J and 76.9% for monkey E.

### Analyses of hand and eye movements

To estimate movement onset, we first calculated the derivatives of the x, y instantaneous hand positions (v_x_ and v_y_ respectively). The two-dimensional velocity was then calculated as:

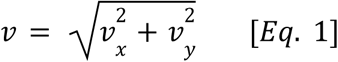

We searched for the moment of peak velocity in the time interval between target onset and target reach. We identified the trough before that peak by finding the moment when the acceleration (second derivative of position) changed sign from negative to positive. We then applied a threshold of 5% * (*v_peak_*− *v_troug_*_ℎ_) to the velocity traces and we defined the movement onset as the first moment after the velocity crosses that threshold. Movement initiation latencies were defined for each hand movement as the difference: moment of movement onset minus moment of target onset (**Figure 2B**). Movement execution durations were calculated as the time elapsed between the movement onset and the moment of target reached. Durations do not necessarily lie in a Gaussian distribution because trajectories do not have identical lengths (**Figure S.1**).

To estimate the proportion of movements where monkeys moved toward the correct target, we computed the angular deviation of the initial hand trajectory from the straight path that connected the previous and the next target (**Figure S.2A** top schema). The initial movement direction was defined by the points A (x_A_,y_A_), with coordinates the x and y position of the hand at the moment of movement onset, and B (x_B_, y_B_), with coordinates the average position of the hand in the first 200 ms after movement onset. The direction of the illuminated target was defined by the previous target position C (x_Tg-1_, y_Tg-1_) and the current target position D (x_Tg_, y_Tg_). The angular deviation from the straight path (in rad) was calculated for each movement:

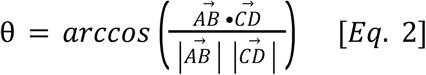

To assess whether monkeys anticipated the upcoming target locations, we computed the cosine similarity between the hand trajectory direction and the target direction (**Figure 2C** top schema). The trajectory direction was defined by the points E (x_t_, y_t_), F (x_t+1_, y_t+1_), whose coordinates are the instantaneous x, y hand positions at time t and t+1, respectively. The target direction was defined by the same point E and the target position D (x_Tg_, y_Tg_). We calculated the cosine similarity at each time point in a window from −300 to +600 ms relative to target onset. In the pre-target period (−300 to 0 ms), hand trajectory direction was calculated from small-amplitude movements performed within the Valid Target Area (1 cm radius) during preparation of the upcoming reach, and target direction corresponded to the upcoming, non-visible target. In the post-target period (0 to +600 ms), target direction refers to the same target after its appearance on the screen. The cosine similarity was computed for each movement j and for each time point t, as the normalized dot product of the two vectors:

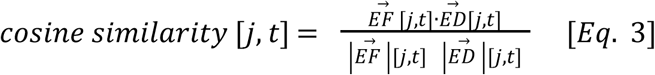

The cosine similarity shows how well the hand movement is aligned to the direction of the target at every instant of every movement. Values of 0 indicate that the two vectors are orthogonal (forming an angle of π/2), values of 1 mean that the two vectors point at the same direction (angle is zero), while values of −1 mean that the two vectors point at the opposite direction (angle is π).

To assess when hand trajectory significantly aligned with target direction across trials, we computed the cosine similarity after shuffling the target direction to one of the possible directions, for each target rank separately (200 shuffles). We tested the null hypothesis that monkeys did not anticipate future target locations. For the first target, monkeys could move from the center towards any direction, so the maximum absolute range of angles they could move was 180°. By averaging many trials, the angle by chance would be around 90° (cosine similarity around 0). Under the null hypothesis, for the second and third targets, monkeys still had to reach one out of six possible targets in the periphery. However, for these targets, the hand was initially located at a peripheral target, and due to workspace geometry subsequent targets would never appear further outward, but rather at one of the other positions in the (invisible) hexagon. Consequently, the range of possible upcoming target positions was restricted and the cosine similarity that can be obtained by chance slightly increased for the second and third targets. We computed the standard deviation of the shuffle distribution across iterations at each time point (**Figure 2C**, shaded intervals). By performing a t-test at each time point, followed by Bonferroni correction for multiple comparisons, we identified significant time periods as temporal clusters in which the observed cosine similarity exceeded the mean plus half standard deviation or fell below the mean minus half standard deviation of the shuffle-derived null distribution.

Decreases in cosine similarity (and negative values) indicate that the hand moved opposite to the target direction. In the pre-target epoch (−300 to 0 ms), this overlapped with the chance distribution (**Figure 2C**) and it related to small adjusting movements within the Valid Target Area, while preparing for the upcoming reach that was often in the opposite direction due to the geometry of sequences (e.g. upward movement was often followed by a downward movement, **Figure S.1**). In the post-target period (0 to +600 ms), similarity decrease in monkey J occurs after the end of the reaches (movement durations were about 200-400 ms, **Figure S.2B**) reflecting the same pre-target movement adjustment but for the next sequential reach.

We computed the tracked eye velocity aligned to the 1^st^, 2^nd^ and 3^rd^ target onsets, for each eye movement j and each timepoint t as the derivatives of the x, y instantaneous tracked eye positions and the two-dimensional eye velocity was calculated as in **Eq. 1**. Generally, the eye signal recordings included a lot of instrumental noise, partly due to the orientation of the horizontal screen below the fixed head of the monkeys. As a result, full trials comprising all three movements were discarded from this analysis when the values of eye velocity were inexplicably high throughout the full trial (or even session) and were unmodulated by the task events. The eye velocities in **Figure 2D** are averaged over the remaining trials, thus the confidence intervals are wider, but the discussed effect is still evident.

To estimate eye movement onset, we applied a velocity threshold of 30 cm/s and defined the onset as the moment at which the eye velocity first crossed this threshold before reaching peak velocity.

In each single trial, we calculated the hand-eye movement initiation delay as the difference between the moment of hand minus eye movement initiation. Positive values indicate that the moment of hand movement onset comes after the moment of eye movement onset, or in other words, the eye moves first and the hand follows. On the other hand, negative values indicate that the hand moves before the eye.

For each behavioral analysis, we performed statistical comparisons between the distributions across target ranks using the non-parametric Wilcoxon signed rank test pairwise (1^st^ – 2^nd^, 1^st^ – 3^rd^, 2^nd^ – 3^rd^ target). We obtained the probability values and we subsequently performed Bonferroni correction for multiple comparisons (3 tests).

### Neural recordings

After training, monkeys were implanted with chronic microelectrode Utah arrays (Blackrock Neurotech) in five cortical areas along the dorsal visuomotor pathway. Four 6×6 channel arrays were implanted in the occipital areas V1,V2 and in the parietal areas Dorsal Prelunate (DP) and Brodmann area 7A, while one 10×10 channel array was implanted in the frontal areas of dorsal premotor and primary motor cortex (PMd/M1). In this study, we focused on the recordings coming from the arrays in area 7A and PMd/M1 (**Figure 3A**).

### Preprocessing

Electrophysiological recordings were acquired at 30 kHz sampling rate. The voltage from each active channel of 7A (n=32) and PMd/M1 (n=96) arrays was bandpass filtered in the range of 600 – 6000 Hz to cut off low frequency oscillations. Activity from single units was isolated using the software KiloSort (Pachitariu et al., 2016) that uses template-matching for spike waveform detection and clustering. Single units with less than 100 spikes in an entire session were excluded from the analyses. Spike trains for each unit were converted to smooth firing rates by convolving them with a one-dimensional gaussian kernel of size 40 ms. The SUA was aligned in epochs from −300 ms until +600 ms around each target onset. A typical recording session yielded about 120 single-units and 25 multi-units across all electrodes in the PMd/M1 array, and about 25 single-units and 15 multi-units in the 7A array.

The envelope of the analog multi-unit activity (MUA) from each channel was computed with the method introduced by Buchwald et al., 1965 and Stark & Abeles, 2007. The root-mean-square of the high-pass filtered signal was computed and was afterwards down-sampled (after using an anti-aliasing zero-phase IIR filter of order 8) at a sampling rate of 1 kHz, matching the behavioral recordings’ sampling rate. Finally, the MUA was z-scored for each channel separately across all time of each session. This mesoscopic neural activity measure has also been described as the spiking-band power and has been shown to improve the decoding performance of brain-machine interfaces (Nason et al., 2020). The MUA for each session was aligned in epochs from −300 ms until +600 ms around each target onset. Trials from all sessions were concatenated, after validating that the variance of each channel recorded across different days was never significantly higher than the variance obtained by chance.

### Assessing anticipatory directional tuning in single-unit activity

To evaluate whether directional tuning of single units emerged earlier for predictable than for unpredictable targets, we first identified for each neuron the directional tuning of the SUA for movements toward the unpredictable 1^st^ target. We then averaged the single-trial gaussian-convolved firing rates in the interval from 100 to 300 ms after the 1^st^ target onset and across all movements in the same direction, for each of the four directions that were repeated in all three target ranks (i.e. top-right, top-left, bottom-right, bottom-left). This yielded the “template” tuning vector of each unit, consisting of one value for each of the four movement directions.

Subsequently, to determine when this template vector re-emerged for movements toward predictable visual targets, we computed the time-resolved Pearson correlation between the template tuning vector and the corresponding four-element SUA vector. To estimate chance level, we recomputed the correlation at each timepoint after randomly permuting the directional labels of the SUA vector while keeping the template tuning vector unchanged, to break the direction-per-direction link between the variables. The chance confidence intervals were then defined as the mean plus/minus one standard deviation across the permutations.

### Assessing initial hand position effects across targets in the sequence

To evaluate whether the observed differences across targets are due to the different hand positions at movement initiation, we trained four Linear Discriminant classifiers to split two classes (**Figure S.4A**). Class 1 comprised the four center-out movements (toward top left, top right, bottom left, bottom right) and class 2 comprised peripheral movements toward the same four directions. Thus classifiers could not separate the two classes based on movement direction. We tested these models on Class 3, which comprised new peripheral movements that were not used during training. In each of the four models, movements included in training class 2 differed. For example, in the first model, training class 2 included movements on the right hand side of the workspace, and test class 3 included movements on the left hand side of the workspace. Accordingly for the remaining three models, we tested all possible combinations of peripheral movements, always comparing them with the center-out movements of training Class 1, after balancing the amount of trials in each class (**Figure S.4A**).

To avoid artifactual classification due to movements that were initiated even before target onset, we trained the classifiers on the preparatory MUA in the 200ms immediately preceding movement onset (MO). We used bins of 50ms starting from MO-200ms with steps of 10ms until MO. Decoding performance was evaluated with respect to how class 3 trials (peripheral movements in the testing set) were labeled by the classifier trained to discriminate class 1 (center-out movements) and class 2 (peripheral movements in the training set). Specifically, the decoding accuracy was calculated as the fraction of correctly predicted movement directions over the total amount of movement directions tested.

To estimate the chance level of the classifiers, we trained the models after having randomly shuffled the labels of the movement directions (200 shuffles). We calculated the decoding accuracy for each shuffle, to create the null distribution of accuracies obtained by chance.

Significantly positive decoding accuracy indicated that class 3 trials were predominantly predicted as class 2 (**Figure S.4B**). This pattern implies that peripheral movements from the testing set occupy a similar region of the latent feature space—defined by their projection onto linear discriminant axes—as peripheral movements from the training set, and are clearly separated from center-out movements. Importantly, this clustering occurred despite differences in initial hand position, indicating that initial position did not provide sufficient discriminative information for the classifier to segregate peripheral movements into distinct groups. This result supports the interpretation that neural activity differences between center-out and peripheral movements cannot be attributed to effects due to different starting positions of the hand.

Significantly negative decoding accuracy indicated that class 3 trials were predominantly predicted as class 1 (**Figure S.4B**). In this case, testing peripheral movements were closer to center-out movements than to training peripheral movements in the discriminant space. Such a pattern reflects a graded influence of initial hand position on the representational geometry of movements in the neural feature space.

Accuracy values within chance-level confidence intervals indicated that class 3 trials could not be reliably associated with either class 1 or class 2, and therefore no representational proximity between these conditions could be inferred.

### Population-level direction decoding using Linear Discriminant Analysis

A hand movement at a given timepoint can be described as a point in a high-dimensional space where each axis/dimension corresponds to the MUA of a single channel for PMd/M1 and another such space for 7A (**Figure 4A**). For visualization purposes, and because the reaching behavior was organized in a two-dimensional workspace, we sought a two-dimensional neural subspace that maximally separated movement directions. To find the dimensions that best separate the movements across directions, we applied a Linear Discriminant Analysis (LDA, Wang et al., 2024). For every movement in each trial, we averaged the MUA in the first 200 ms after the 1^st^ target onset, a window that approximates the planning epoch of the movement towards an unpredictable target. We trained two LDA models (one for PMd/M1 and one for 7A) to maximally separate the movement directions based on the provided labels of the actual trajectory directions.

Each center-out hand movement had one out of six possible specific movement directions. However, due to the geometry of the sequences (**Figure S.1**), only movements toward the top-right, top-left, bottom-right and bottom-left were common to all target ranks (1^st^, 2^nd^ and 3^rd^). Horizontal leftward and rightward movements occurred only for the 1st target, and vertical upward and downward movements occurred only for the 2nd and 3rd targets.

After identifying the discriminant axes from the six center-out directions of the 1^st^ target, we projected the MUA of the movements toward the 2^nd^ and 3^rd^ targets onto the same axes (four common directions in **Figure 4B**, all directions in **Figure S.5A**, 7A projections in **Figure S.5B**). For movements toward the 1^st^ target, we applied cross-validation by training the LDA on 80% of the trials and projecting only the remaining 20%.

To assess whether a common directional representation was present across target ranks and emerged earlier for movements toward the 2nd and 3rd targets, we trained LDA classifiers (one per area) to discriminate across the four directions common to all target ranks (i.e. top-right, top-left, bottom-right, bottom-left). Horizontal leftward and rightward movements occurred only for the 1st target and were therefore excluded from the training set, as they could otherwise be incorrectly predicted for the 2nd and 3rd targets, artificially reducing decoding accuracy since these labels do not exist in the test set. Likewise, vertical upward and downward movements could not be assigned to any class in the classifier trained on the 1st target. We then evaluated whether, and at which time points, the classifiers could predict the target direction of movements toward the 1^st^ target (cross-validated as previously), 2^nd^ and 3^rd^ targets. Decoding accuracy was defined as the number of correctly classified movements divided by the total number of tested movements (**Figure 4D**).

We tested the predictions against the null hypothesis that MUA directional tuning during the second and third targets differed from that observed during the first target. Under this hypothesis, a classifier trained to predict target direction from MUA activity during the first target should not generalize to the second and third targets and therefore should fail to predict target direction above chance level.

To estimate the chance level, we trained LDA classifiers after randomly shuffling the labels of the movement directions across trials (200 shuffles). We computed the decoding accuracy for each shuffle to generate a null distribution of accuracies expected by chance. Because the classifier predicted one of four possible target directions, chance performance was approximately 25% for all target ranks.

Decoding accuracy at chance level indicates that the directional information encoded in MUA during the tested condition does not generalize from the training condition, preventing accurate prediction of target direction. Conversely, decoding accuracy significantly above chance indicates that directional information was preserved across target ranks, such that similar neural activity patterns are associated with the same movement directions.

### Identifying channels with significant directional modulations

To identify channels in which the MUA was significantly modulated by movement direction, we computed the Mutual Information (MI) between the MUA and the discrete variable of instructed movement direction. To compute the MI and the null hypothesis, obtained by calculating the MI after shuffling the movement direction labels (200 shuffles), we used the python package Frites (Combrisson et al., 2022). For each target rank separately, we computed the MI and the clusters of significant time points corrected for multiple comparisons using cluster-based statistics.

Channels were considered directionally modulated if they exhibited at least one cluster of consecutive significant time points spanning 100 ms. Under this criterion, 20 of 32 (62.5%) channels in 7A and 68 of 96 (70.8%) channels in PMd/M1 were classified as significant in monkey J, compared with 22 of 32 (68.8%) and 77 of 96 (80.2%) channels, respectively, in monkey E.

### Feature-specific Information Transfer and motor-parietal interactions

To assess the directed inter-areal interactions between each pair of channels, we calculated the Feature-specific Information Transfer (FIT; Celotto et al., 2023). This information theoretic measure quantifies the amount of information-related to a specific task parameter-that is transferred between two regions, similar to Transfer Entropy and Wiener-Granger Causality measures (Granger, 1969). The advantage of the FIT when compared to the aforementioned metrics is the task-related specificity of the information that is transferred. The FIT is included in and bounded by the Transfer Entropy between the sender and the recipient and is measured in bits. Extensive proofs and validations of this measure on simulations and real datasets are presented in Celotto et al., 2023.

We calculated the FIT about the movement direction for all pairs of channels (i.e. 6144 pairs) between 7A and PMd/M1 in windows of 200 ms aligned on each target onset. The combination of channel pairs between 7A and PMd/M1 in which both channels exhibited significant MI was calculated as:

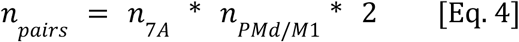

The product of the channels is multiplied by 2 to account for both directions (i.e. 7A → PMd/M1 and PMd/M1 → 7A). There were 2720 out of 6144 (44.2%) channel pairs in monkey J and 3388 out of 6144 (55.1%) pairs in monkey E that included one channel from 7A and one from PMd/M1 with significant directional modulations.

We found 874 pairs with significant FIT in monkey J and 332 in monkey E. This corresponds to 14.2% out of 6144 possible pairs or 32.1% out of 2720 modulated pairs in monkey J and 5.4% out of 6144 possible pairs or 9.8% out of 3388 modulated pairs in monkey E.

A maximum lag of 100 ms was used for the calculation of the past activity. The FIT for each channel pair was computed for each lag separately, resulting in a 3D matrix of amplitude, lag, time, and it was then aggregated across lags. The FIT increased when information about the movement direction in the MUA from a source channel predicted the information about direction in the present activity from a target channel (**Figure 5A**). Importantly, the FIT captured only the additional information that was present in the past of the source that did not already exist in the past of the target itself.

To assess the significance of FIT, we applied non-parametric cluster-based statistics. More specifically, we disrupted the link between the MUA and the respective trajectory direction of that movement, by shuffling the labels of the direction (200 shuffles) for each individual pair of channels between 7A and PMd/M1. We computed the FIT for each surrogate to obtain the null distribution of information values and we set the cluster forming threshold at the 95^th^ percentile of the null distribution. The p-values (corrected for multiple comparisons) were then calculated as the proportion of cluster masses maxima in the null distribution that surpassed the cluster mass maximum of the true FIT (Combrisson et al., 2022). We considered a pair of channels significant only if it was surpassing the significance threshold (i.e. p < 0.05) for at least 50 ms consecutively and we sorted the significant channel pairs into different groups (**Figure 5B, Figure S.9**). The bottom group consisted of pairs of channels having a bidirectional information flow between the two areas. The middle group included pairs that showed significant information flow uniquely PMd/M1 → 7A, while the top group included only pairs that were significant in the direction 7A → PMd/M1. The rest channel pairs did not pass the significance thresholds for any of the two directions.

To assess whether channels with significant FIT were spatially localized or distributed in the arrays, we computed the net FIT (**Figure S.8A**). To calculate it we subtracted the total information that each PMd channel “sends” to all 7A channels-outward connectivity-from the total information that each PMd channel “receives” from all 7A channels-inward connectivity, as follows:

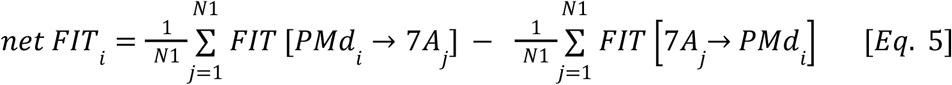

where N1 is the amount of 7A channels (i.e. 32). For each channel of the 7A array the net FIT was found by subtracting “outward” from “inward” (opposite than for PMd/M1 to maintain the same sign of the net FIT), as follows:

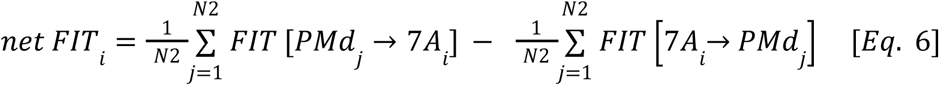

where N2 is the amount of PMd/M1 channels (i.e. 96). Thus, in both arrays, positive values of net FIT indicate an asymmetry in the PMd/M1 → 7A direction, while negative values show an asymmetry in the opposite direction (7A → PMd/M1).

To examine whether channels with significant FIT were localized or distributed across each array, we calculated the spatial autocorrelation by computing the square difference between the net FIT of each channel and the average net FIT of its neighboring channels, up to the 5th nearest neighbor (**Figure S.8B**). To facilitate the comparisons, we normalized the square difference by the maximum square difference that could possibly exist in each array. The mean squared difference obtained by chance was defined by recomputing the difference after shuffling the channel labels.

For each channel pair separately, we identified the FIT peak as the maximum value in the three-dimensional space defined by amplitude, latency, and lag. This yielded, for each pair in each direction, a specific amplitude, latency, and interaction lag at which information transfer was maximal (**Figure 5C, Figure S.10**). The average FIT profile for the direction PMd/M1 → 7A (**Figure 5D**) was calculated by averaging the FIT across all pairs of the middle group (**Figure 5B**), while the FIT for the opposite direction (7A → PMd/M1) was averaged across all pairs of the top group (**Figure 5B**).

Within the FIT framework, we cannot exclude that, even though the amount of information about direction in the past might be predictive of the future, it could arise from different neural representations. To examine whether the representations are the same, we used Canonical Correlation Analysis (CCA). We identified a pair of dimensions (one for PMd/M1 and one for 7A, **Figure S.7A**) that captures their covariance across the four movement directions and maximizes the projection correlation between the two areas (Semedo et al., 2019). We applied regularization to ensure that only the variance, and not the amplitude of the activity, would affect the correlation estimation. To ensure the convergence to a stable result, we applied the CCA on the channels with peak MUA amplitude above the 75^th^ percentile in each array separately. We performed a new CCA for each timepoint aligned around the 1^st^, 2^nd^ and 3^rd^ target onsets, separately. The covariance matrix was computed in bins of 50 ms, but similar results were obtained with bins of 10 and 100 ms. High correlation indicated that trials shared a similar representational ordering of direction across the two areas, whereas low correlation reflected the absence of such structure (**Figure S.7B**). To compare the correlation differences across the three targets, we resampled the projected trials (bootstrap with replacement) and computed multiple projection correlations (20 bootstraps). We obtained the final correlation by averaging over the bootstraps and the confidence intervals represented the 95^th^ percentile of the distribution of the bootstraps (**Figure S.7C**). Finally, to estimate the null correlation by chance, we calculated the CCA after randomly shuffling the trials for the PMd/M1 MUA, breaking the trial-to-trial relationship between PMd/M1 and 7A.

### Permutation-based statistics

To calculate the significance of differences across target ranks (1^st^, 2^nd^, 3^rd^ in **Figure S.3** and **Figure S.10**), we used permutation testing. This is a non-parametric statistical test that makes no assumptions on the distribution of the data and is especially useful when we compare groups of dependent, but unpaired, observations with unequal sizes. For two distributions A and B, we computed the absolute difference of their means. Then, we pooled them together into one distribution AB, we permuted the observations randomly (10000 permutations) and we split them into two new distributions A’ and B’ with the respective sizes of A and B. For each permutation, we computed again the absolute difference between A’ and B’ to get the null distribution. The p-values were calculated as the proportion of the null distribution that was larger than the real difference observed from the data. Bonferroni correction for multiple comparisons was applied, by multiplying the p-values by 3, since we performed three tests: 1^st^ – 2^nd^, 1^st^ – 3^rd^, 2^nd^ – 3^rd^ targets.

## Acknowledgments

We would like to thank Etienne Combrisson, Bjørg Kilavik, Simon Nougaret, Laura López Galdo and Nilanjana Nandi for many discussions and feedback on this work. We also acknowledge the Mediterranean Primate Research Center for the dedicated animal care.

## Ethics statement

All experimental procedures were approved by the local ethical committee (C2EA 71; authorization Apafis#13894-2018030217116218v4) and conformed to the European and French government regulations.

## Author Contributions

Funding acquisition: T.B., A.B., S.G., A.R.; Experimental task design: T.B., S.G., A.R., F.B.; Experiments: S.J., F.B., T.B.; Data analysis: G.B.; Writing - original draft: G.B.; Writing - review & editing: G.B., T.B., A.B., N.M., A.R.; Visualization: G.B., T.B., A.B.

## Data Availability

Raw or preprocessed data can be shared upon reasonable request to Thomas Brochier or Sonja Grün.

## Code Availability

Source code used in this study is available at the public Github repository https://github.com/YorgosBardanikas/vision4action.

