## Supplementary Figures for "Reciprocal Fronto-Parietal Interactions Support Distinct Motor Strategies during Sequential Reaching"

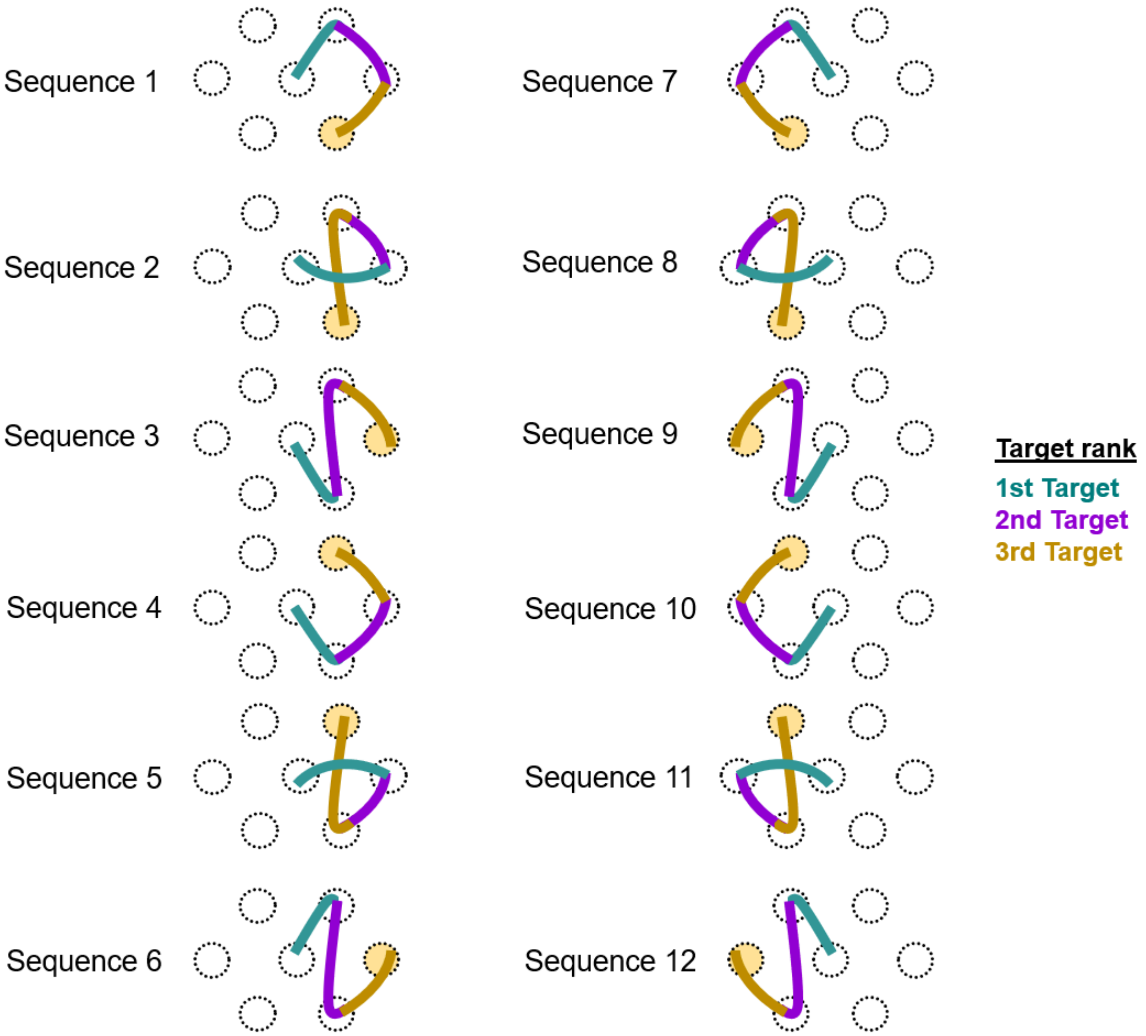


**Figure S.1. The 12 three-target sequences.** Sequences 1-3 are also shown in **Figure 1C**. Sequences 4-6 are the sequences 1-3 flipped vertically. Accordingly, sequences 6-12 are the sequences 1-6 flipped horizontally. To initiate a trial-sequence, monkeys had to dwell for 150 ms in the central target. The last target of each sequence is highlighted with yellow. Monkeys could only see one active target throughout the trial.

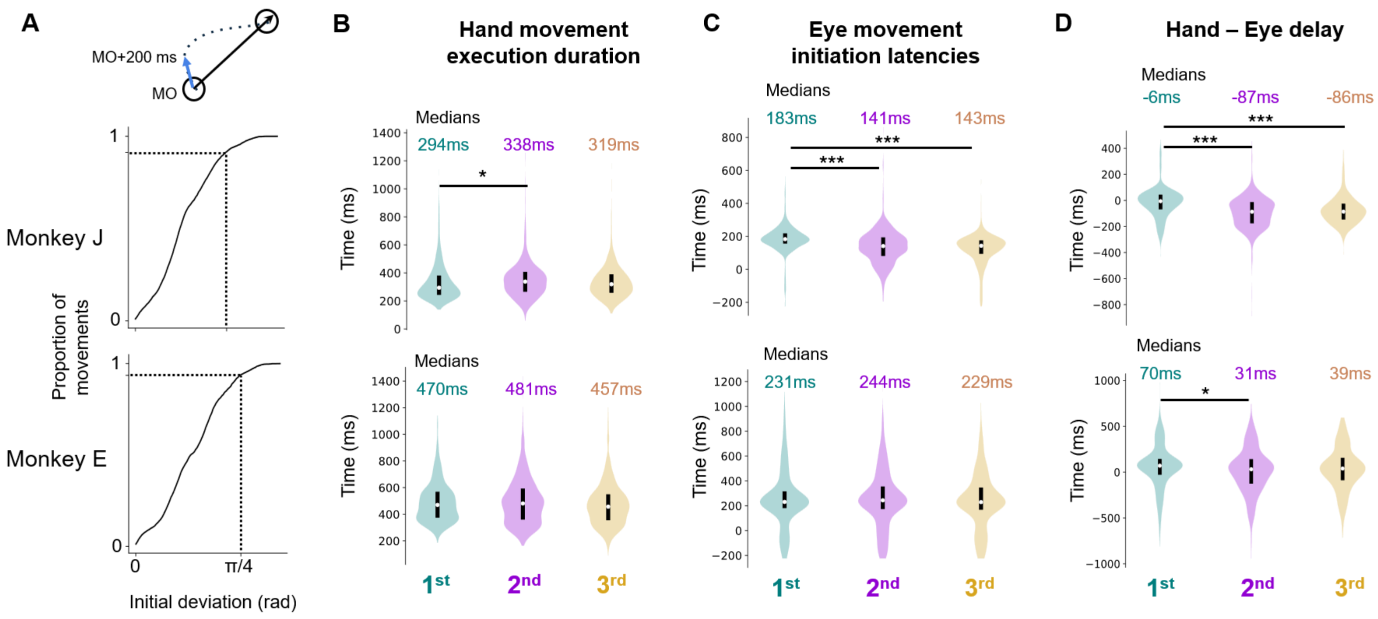


**Figure S.2. Distributions of relevant behavioral variables.** (A) Cumulative sum of the angular deviation between the initial hand trajectory -based on the first 200 ms after the movement onset- (blue vector on top schema) and the straight path between previous and next target (black vector on top schema). Dashed lines show the percentage of movements (from all sessions) whose deviation was smaller than π/4 rad (45°). (B) Distributions of duration of hand movement execution for each target rank. Bimodalities are due to different trajectory lengths. Distribution medians are shown to indicate the effect size across targets. (C) Distributions of eye movement initiation latencies for each target rank. (D) Distributions of single-trial temporal delays between the hand and eye movement initiation (Hand - Eye). Positive values show that hand movement initiation occurs after the eye movement initiation, while negative values show that hand movement occurs first and eye movement follows. Solid black lines between the violin plots indicate the significant differences between the distribution means. P values were computed applying Wilcoxon signed rank tests with Bonferroni correction: $p < 10^{-2}$ is noted with * ; $p < 10^{-20}$ is noted with ***.

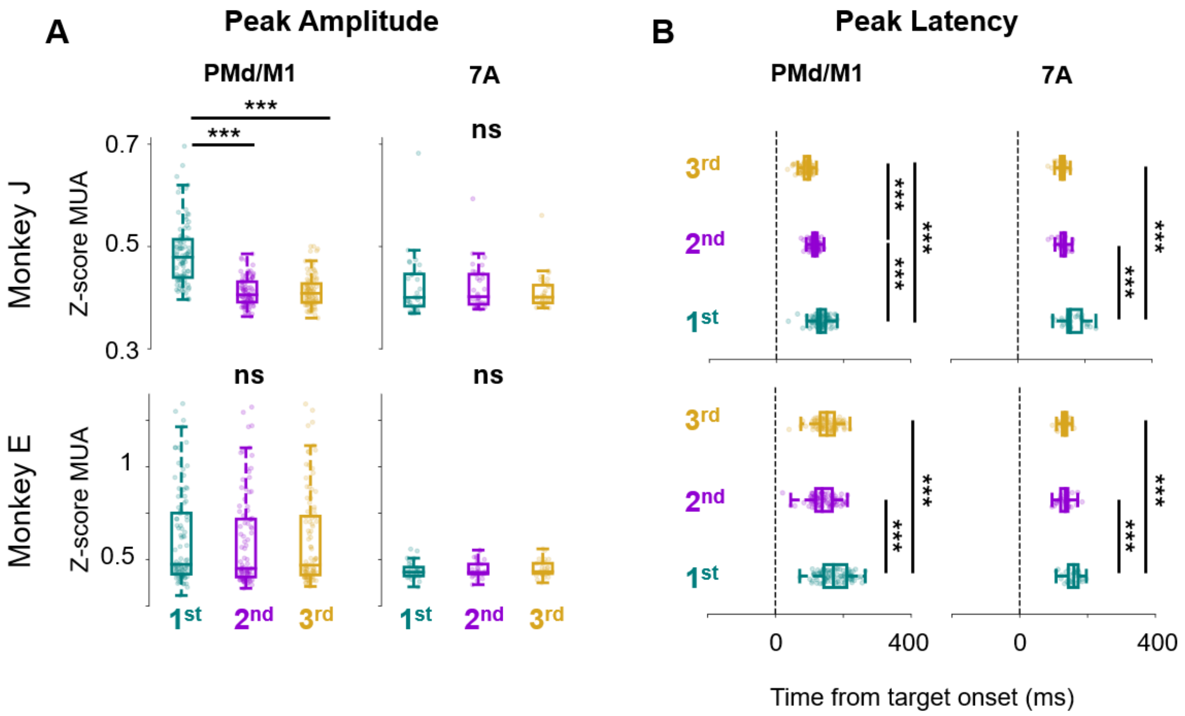


**Figure S.3. Statistics of MUAe peak amplitude and latency across targets.** (A) Distributions of the amplitude of the first MUA peak across channels. Each dot represents the mean peak amplitude of one channel (n=96 for each boxplot of PMd/M1 and n=32 for each boxplot of 7A), averaged over trials in each target condition. The peak amplitude was calculated in each single trial and then averaged across trials for each channel. (B) Distributions of the latency of the first MUA peak across channels. Each dot represents the mean peak latency of one channel, averaged over trials in each target condition. Solid black lines indicate the significant differences between the distribution means P values were computed from permutation testing after Bonferroni correction: $p < 0.05$ is noted with * ; $p \approx 0$ is noted with ***.

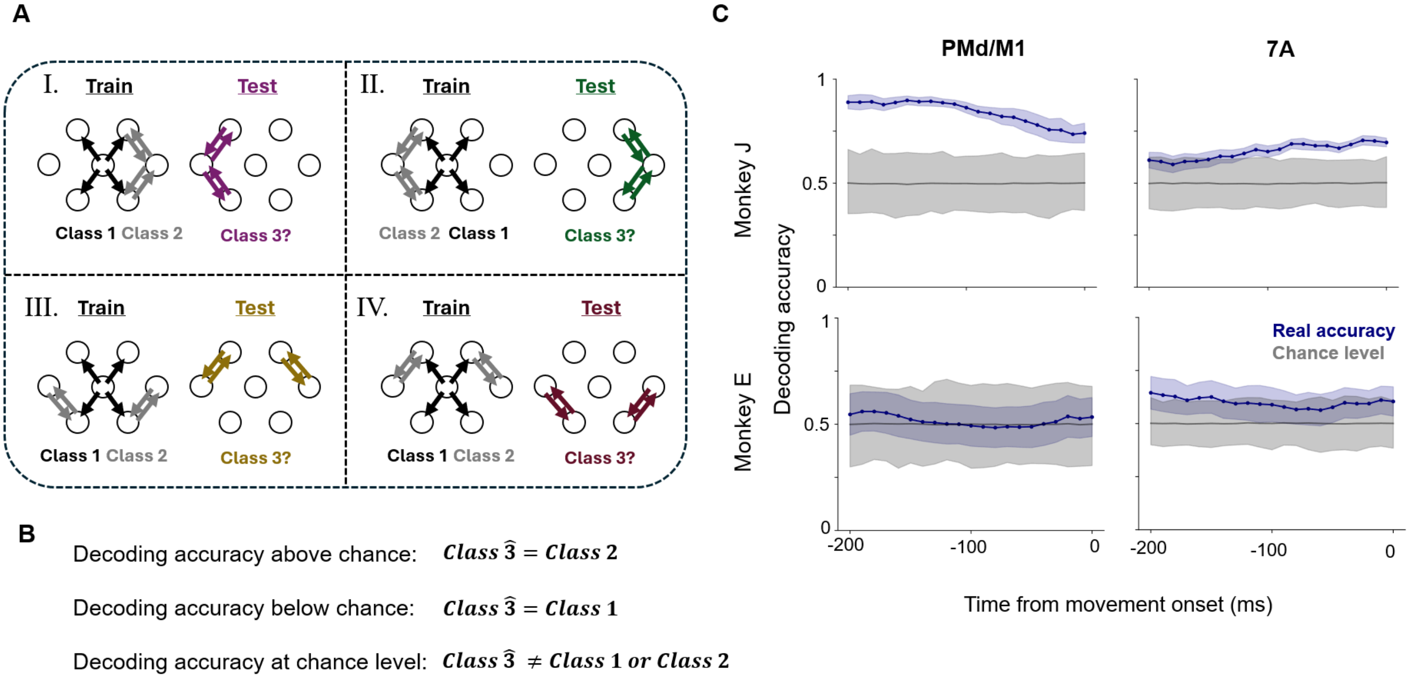


**Figure S.4. PMd/M1 activity differences across targets cannot be explained by different hand positions at movement initiation.** (A) We trained 4 classifiers to split two classes: the center-out movements (across all directions, Class 1) from peripheral movements (across all directions, Class 2) and we tested these models on new peripheral movements (across all directions, Class 3). Class 1 always comprised the center-out movements (black), while class 2 (grey) differed in each of the 4 tests (e.g. in test I., class 2 included movements on the right hand side and we tested movements on the left hand side -class 3). (B) If the decoding accuracy is above chance level (defined by 200 models trained on shuffled labels), class 3 movements are predicted as class 2, whereas if it is below chance class 3 is predicted as class 1. If the accuracy falls into the chance level confidence intervals, then class 3 cannot be associated with any of the other classes. (C) The decoding accuracy of the predicted Class 3 (blue) averaged across the four tests shown in (A). The shaded grey area represents the 95th percentile of chance level and the grey line is the average (centered at 0.5, due to the existence of 2 classes). The accuracy in monkey J was above chance, so the neural activity during all peripheral movements shared commonalities among them, despite their complete opposite initial hand positions, and differences from the center-out movements. Thus, differences in MUAe across target conditions for monkey J (observed in Figure 3**C**) could not be explained by different initial hand positions before movement initiation and were more likely related to the different predictability levels between center-out and peripheral movements.

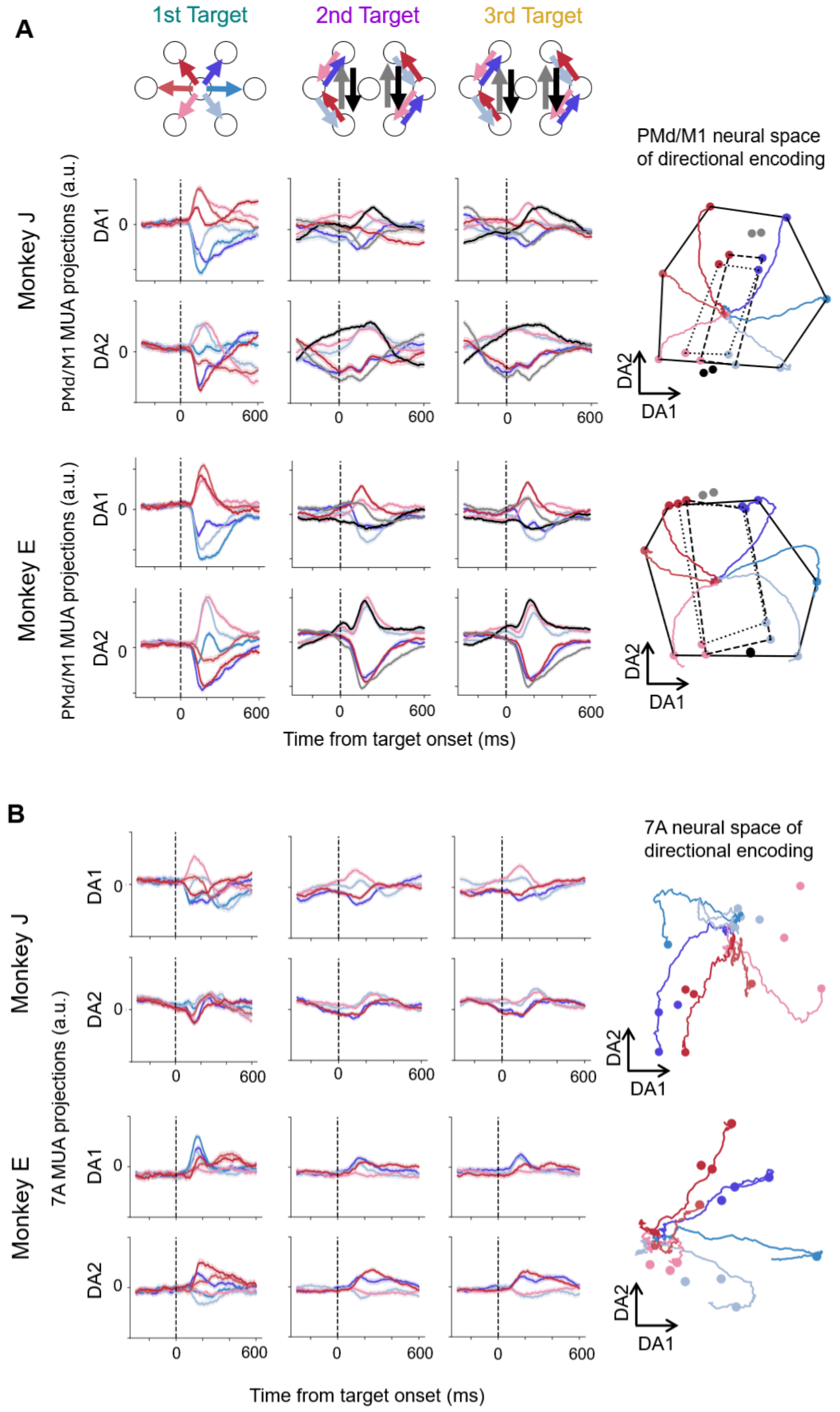

A
1st Target
2nd Target
3rd Target
PMd/M1 neural space of directional encoding
Monkey J
Monkey E
PMd/M1 MUA projections (a.u.)
DA1
DA2
0
600
Time from target onset (ms)
B
7A neural space of directional encoding
7A MUA projections (a.u.)

**Figure S.5. Neural representations of directional tuning in 7A and PMd/M1.** (A) Right. PMd/M1 MUA projections including vertical movements toward the top (grey lines), and toward the bottom (black lines). Each of the two rows shows one discriminant axis (DA1 and DA2), while each of the three columns indicates the epoch around the 1st , 2nd and 3rd target respectively. Each line shows the average across movements to the same direction in arbitrary units. Confidence intervals show the 95th percentile of the estimation of the mean (200 bootstraps). Left. The two-dimensional neural space composed of the two discriminant axes. The colored dots (one per direction) show the projection at the moment of the maximum variance across directions for each target rank. Vertical top movements (grey dots) are projected between the top-right and top-left, while vertical bottom movements (black dots) between bottom-right and bottom-left accordingly. (B) Same as in (A) but for the 7A projections on the LDA axes built from 7A neural activity (0-200ms after 1st target onset). The two-dimensional 7A neural space was not organized in the same hexagonal manifold as in PMd/M1, however movements toward the same direction were still clustered together, which allowed above chance decoding of future direction in **Figure 4D**.

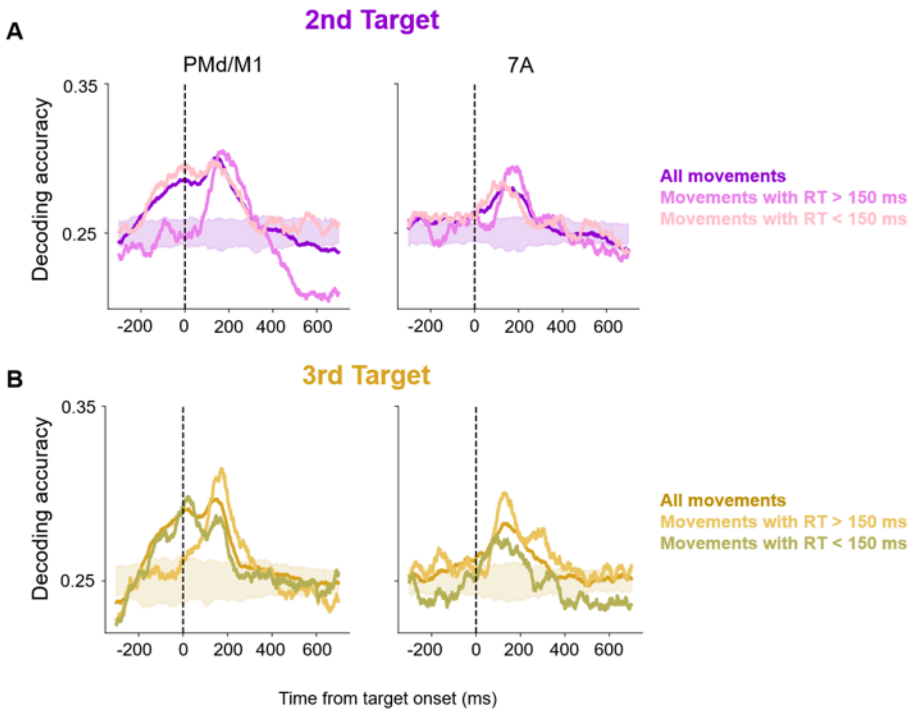


**Figure S.6. LDA accuracy in function of reaction times.** Decoding accuracy of movement direction for monkey J, who generally anticipated movements to predictable 2nd (A) and 3rd (B) targets. Decoding accuracy of movement direction for all movements, shown in **Figure 4D**, is compared to the accuracy of movements with reaction times shorter or longer than 150 ms (typical reaction time of macaque monkeys). Movements in these two conditions have been randomly resampled to avoid accuracy biases due to imbalanced samples. The shaded intervals show the distribution of accuracy by chance obtained by LDA models trained on shuffled direction labels. Movements with long reaction times reflect a proxy of absence of motor anticipation and they do not cluster with the rest; in contrast they become significant after the predictable target onset.

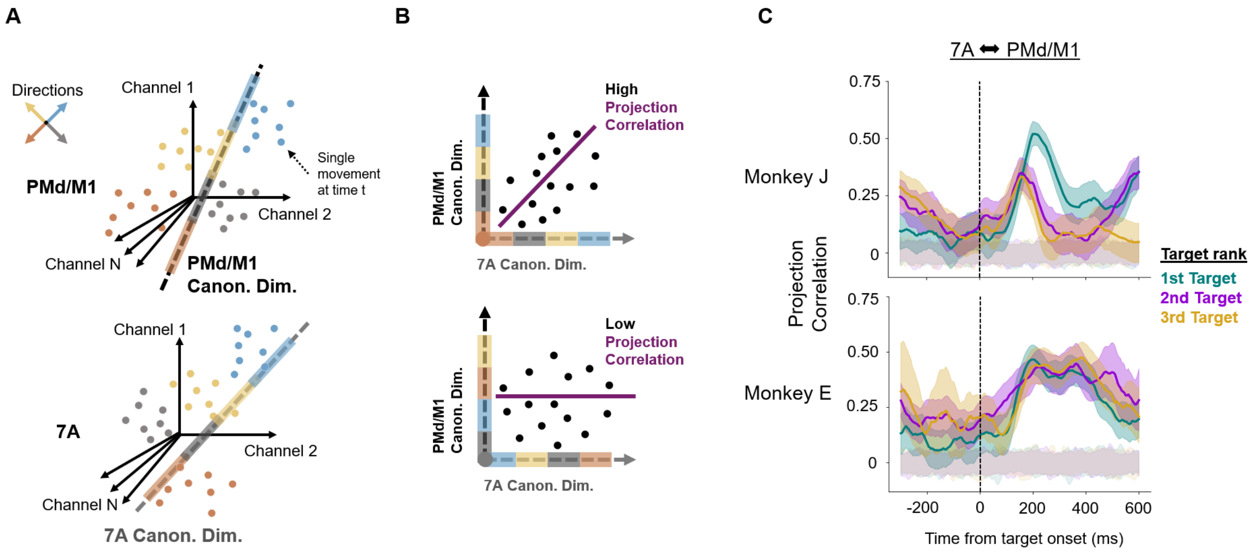


**Figure S.7. Covariance across movement directions between 7A and PMd/M1.** (A) We applied Canonical Correlation Analysis (CCA) to identify a pair of dimensions (one in the PMd/M1 space and one in the 7A space) that captures the maximum covariance across movement directions between the two areas. In principle, trials from the same direction cluster closely, while trials from different directions are farther apart. (B) The projection correlation across trials (purple line) is high when direction-based clusters of projected trials follow a similar ordering in both areas, and low when this correspondence is absent. (C) The projection correlation between PMd/M1 and 7A across the epochs around the 1$^{st}$ (teal), 2$^{nd}$ (violet) and 3$^{rd}$ (gold) targets. Confidence intervals show the 95th percentile of the estimation of the mean correlation (200 bootstraps). The null correlation (shaded intervals) was computed for each timepoint and each segment from models of shuffled trials. Significant correlations (values surpassing the null) show that the ordering of the four directions is the same in the neural representations of both PMd/M1 and 7A.

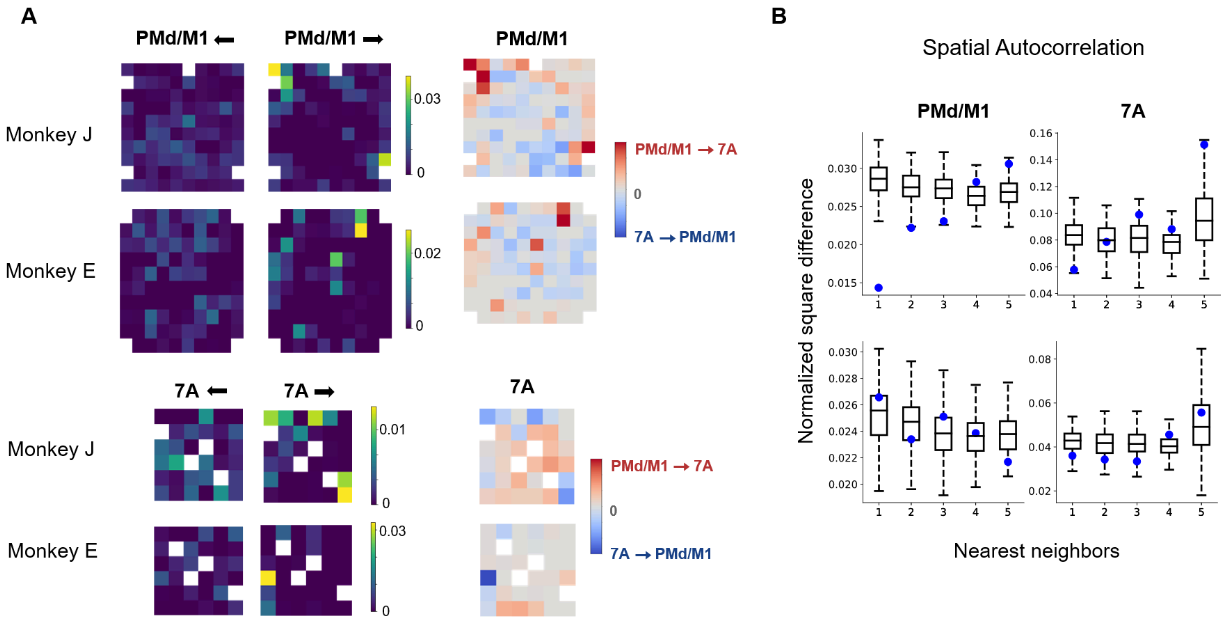


**Figure S.8. Distribution of FIT in the PMd/M1 and 7A channels.** (A) Left. Inward (⟵) and outward (⟶) connectivity for each PMd/M1 and 7A channel. The channel per channel subtraction of the two directions leads to the map of net FIT in the right. Right. Net FIT about the movement direction around the 1st target onset for each channel of each array. Warm colors indicate asymmetric FIT in the direction PMd/M1 ⟶ 7A while cold colors indicate an asymmetry in the opposite direction (7A ⟶ PMd/M1). (B) Normalized square difference between the net FIT of each channel with its neighboring channels (up to the 5th nearest neighbor) for each array. Blue dots represent the true effect while the boxplots show the distribution of the square difference after shuffling the channels, reflecting the null hypothesis (difference due to chance).

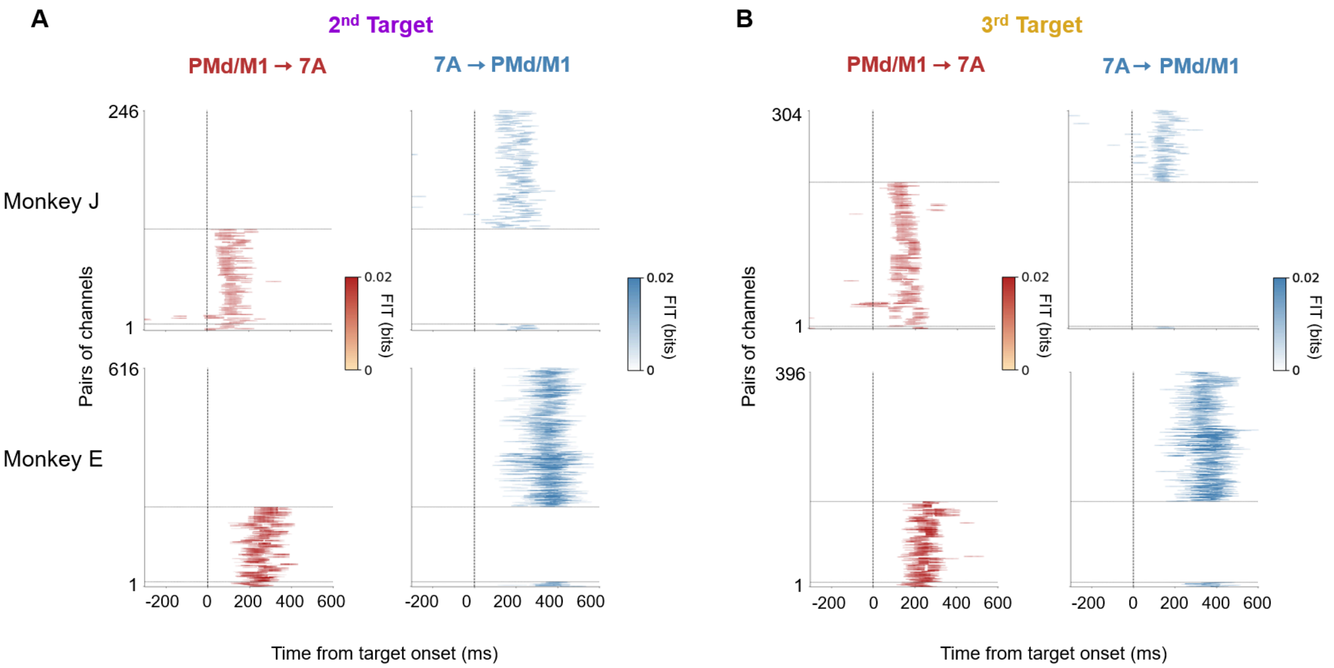


**Figure S.9. Significant FIT clusters.** Similar to **Figure 5B**. The FIT clusters for all significant channel pairs around the (A) 2nd target onset and (B) 3rd target onset.

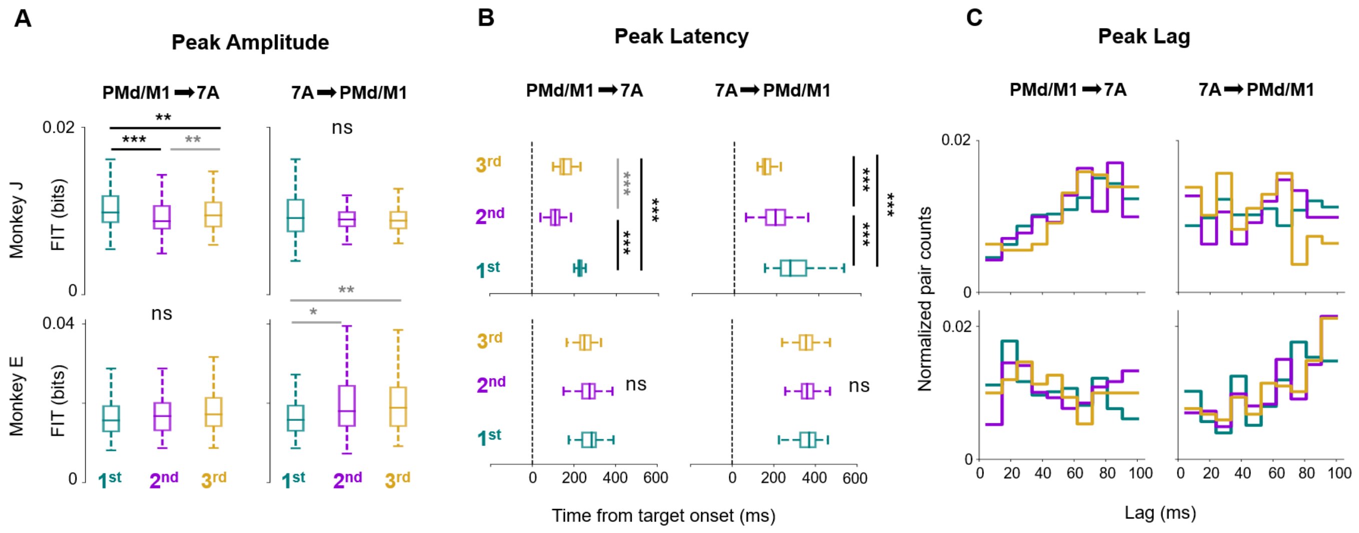


**Figure S.10. Statistics of FIT peak across significant channel pairs.** (A) Distribution of amplitudes of the FIT peak for all significant pairs of channels for the three target ranks in each direction. Solid lines indicate the significant differences between the distribution means. Black lines indicate that the first distribution mean is significantly larger than the second. While grey lines mark the opposite. P values were computed from permutation testing after Bonferroni correction: p < 0.05 is noted with * ; p ≈ 0 is noted with ***. (B) Distribution of latencies of the FIT peak for all significant pairs of channels for the three target ranks in each direction. (C) Distribution of lags of the significant FIT peak for channel pairs across targets. The pair counts were kernel-normalized so the area under the curve integrates to 1. Kolmogorov-Smirnov pairwise tests across targets could not reject the null hypothesis that the lag distributions across targets were sampled from the same underlying distribution.
